# ACDC: Automated Cell Detection and Counting for Time-lapse Fluorescence Microscopy

**DOI:** 10.1101/2020.07.14.202804

**Authors:** Leonardo Rundo, Andrea Tangherloni, Darren R. Tyson, Riccardo Betta, Carmelo Militello, Simone Spolaor, Marco S. Nobile, Daniela Besozzi, Alexander L. R. Lubbock, Vito Quaranta, Giancarlo Mauri, Carlos F. Lopez, Paolo Cazzaniga

## Abstract

Advances in microscopy imaging technologies have enabled the visualization of live-cell dynamic processes using time-lapse microscopy imaging. However, modern methods exhibit several limitations related to the training phases and to time constraints, hindering their application in the laboratory practice. In this work, we present a novel method, named Automated Cell Detection and Counting (ACDC), designed for activity detection of fluorescent labeled cell nuclei in time-lapse microscopy. ACDC overcomes the limitations of the literature methods, by first applying bilateral filtering on the original image to smooth the input cell images while preserving edge sharpness, and then by exploiting the watershed transform and morphological filtering. Moreover, ACDC represents a feasible solution for the laboratory practice, as it can leverage multi-core architectures in computer clusters to efficiently handle large-scale imaging datasets. Indeed, our Parent-Workers implementation of ACDC allows to obtain up to a 3.7× speed-up compared to the sequential counterpart. ACDC was tested on two distinct cell imaging datasets to assess its accuracy and effectiveness on images with different characteristics. We achieved an accurate cell-count and nuclei segmentation without relying on large-scale annotated datasets, a result confirmed by the average Dice Similarity Coefficients of 76.84 and 88.64 and the Pearson coefficients of 0.99 and 0.96, calculated against the manual cell counting, on the two tested datasets.

## 1. Introduction

Advances in microscopy imaging technologies have enabled the visualization of dynamic live-cell processes using time-lapse microscopy methodologies [1–3]. Therefore, collections of microscopy images have become a primary source of data to unravel the complex mechanisms and functions of living cells [4]. The vast quantity and complexity of the data generated by modern visualization techniques preclude visual or manual analysis, therefore requiring computational methods to infer biological knowledge from large datasets [5,6]. In particular, the objects of interest in cellular images are characterized by high variations of morphology and intensity from image to image [6], which make features such as cell boundaries and intracellular features difficult to accurately identify. For this reason, cell segmentation analysis has gained increasing attention over the last decade [7].

The most commonly used free and open-source software tools for microscopy applications in the laboratory are ImageJ [8] or Fiji [9], and CellProfiler [10]. Although these tools offer customization capabilities, they do not provide suitable functionalities for fast and efficient high-throughput cell image analysis on large-scale datasets. In addition, CellProfiler Analyst [11] allows the user to explore and visualize image-based data and to classify complex biological phenotypes with classic supervised Machine Learning (e.g., Random Forests, Support Vector Machines). Taken together, these tools provide accurate performance for image quantification on high-quality annotated datasets but generally lack capabilities to work in the laboratory practice, because training and model setup phases are required; in addition, the user is often forced to transfer data from one tool to another for achieving the desired analysis outcome. Therefore, this hinders these tools to match the time constraints imposed by time-lapse microscopy studies.

Various mathematical morphology methodologies have been extensively used in cell imaging to tackle the problem of cell segmentation [7]. Wählby *et al*. [12] presented a region-based segmentation approach in which both foreground and background seeds are used as starting points for the watershed segmentation of the gradient magnitude image. Since more than one seed could be assigned to a single object, initial over-segmentation might occur. In [13], the authors developed an automated approach based on the Voronoi tessellation [14] built from the centers of mass of the cell nuclei, to estimate morphological features of epithelial cells. Similarly to the approach presented in this paper, the authors leveraged the watershed algorithm to correctly segment cell nuclei. It is worth noting that we do not exploit Voronoi tessellation since our objective disregards the morphological properties of adjacent cells in a tissue. Kostrykin *et al*. [15] introduced an approach based on globally optimal models for cell nuclei segmentation, which exploits both shape and intensity information of fluorescence microscopy images. This globally optimal model-based approach relies on convex level set energies and parameterized elliptical shape priors. Differently, no detection with ellipse fitting was exploited in [16], as this shape prior is not always suitable for representing the shape of the cell nuclei because of the highly variable appearance. In particular, a two-stage method combining the split-and-merge and watershed algorithms was proposed. In the first splitting stage, the method identifies the clusters by using inherent characteristics of the cell (such as size and convexity) and separates them by watershed. The second merging stage aims at detecting the over-segmented regions according to the area and eccentricity of the segmented regions. These sub-divisions are eliminated by morphological closing.

Machine Learning techniques have been applied to bioimage analysis [17,18]. For instance, CellCognition aims at annotating complex cellular dynamics in live-cell microscopic movies [19] by combining Support Vector Machines with hidden Markov models to evaluate the progression by using morphologically distinct biological states. More recently, it has been shown that methods based on Deep Convolutional Neural Networks (DCNNs) [20,21] can successfully address detection and segmentation problems otherwise difficult to solve by exploiting traditional image processing methods [22]. Further, an ensemble of DCNNs defined to segment cell images was presented in [23], where a gating network automatically divides the input image into several sub-problems and assigns them to specialized networks, allowing for a more efficient learning with respect to a single DCNN.

With regard to the most popular architectures for object detection and instance segmentation, such as Faster Region-CNN (R-CNN) [24] and You Only Look Once (YOLO) [25], these approaches require a considerable amount of labeled images and yield only a coarse-grained segmentation in cell imaging [26] and medical image analysis [27]. This does not ensure a precise separation between adjacent cells. Even though transfer learning (i.e., the use of pre-trained CNNs on large-scale datasets of natural images) could be applied, hundreds of accurately annotated input samples should be available [28]. Therefore, parameter-efficient architectures, including simple trainable activation functions [29] or mixed-scale dense CNNs [30], might be beneficial to deal with the paucity of manually labeled and validated datasets. Alternatively, also data augmentation techniques based on Generative Adversarial Networks (GANs) [31,32] or interactive solutions [33], require the time-consuming annotation by the experts.

Traditional image segmentation approaches often require experiment-specific parameter tuning, while DCNNs require large amounts of high-quality annotated samples or ground truth. The ground truth, representing the extent to which an object is actually present, is usually delineated by a domain expert, *via* a tedious, cumbersome and time-consuming visual and manual approach. The annotated samples for one dataset may not be useful for another dataset, so new ground truth generation may be needed for a new dataset, thus limiting the effectiveness of DCNNs. The authors of [22] proposed an approach for automatically creating high-quality experiment-specific ground truth for segmentation of bright-field images of cultured cells based on end-point fluorescent staining, then exploited to train a DCNN [34]. In general, applying DCNNs to microscopy images is still challenging due to the lack of large datasets labeled at the single cell level [34]; moreover, Gamarra *at al*. [16] showed that watershed-based methods can achieve performance comparable to DCNN-based approaches. Thus, unsupervised techniques that do not require a training phase (i.e., data fitting or modeling) represent valuable solutions in this practical context [19,35].

In this work, we present a novel method named Automated Cell Detection and Counting (ACDC), designed for time-lapse microscopy activity detection of fluorescent-labeled cell nuclei. ACDC is capable of overcoming the practical limitations of the literature approaches, mainly related to the training phases or time constraints, making it a feasible solution for the laboratory practice. Recent developments of automated microscopy imaging systems has allowed for the generation of very large datasets [36], which are typically ~ 3TB, including over 300,000 images, for each experiment. Each dataset produces novel features due to differences in the cells used, their fluorescence label intensity, and their specific responses to anti-cancer treatment conditions. This variation poses challenges to using deep learning-based segmentation [37] that is trained without including the new data: for any comparisons across datasets, the segmentation would need to be repeated/updated to include every new dataset and re-applied to all previous data. To cope with these issues, ACDC uses bilateral filtering [38] applied on the original image to smooth the input cell image while preserving edge sharpness. This is followed by watershed transform [39,40] and morphological filtering [41,42], applied in a fully automatic manner. Thus, ACDC efficiently yields reliable results by only requiring the settings of a few parameters, which may be conveniently adjusted by the user. Therefore, unlike sophisticated solutions that do not provide any interactive control, ACDC makes the end-user sufficiently aware of the underlying automated analysis, thanks to the resulting interpretability of the segmentation model. We demonstrate applications of ACDC on two different cell imaging datasets in order to show its reliability in different experimental conditions.

The main contributions of this work are summarized hereafter:

- ACDC is a fully automatic pipeline for cell detection and counting that exploits watershed transform [39,40], morphological filtering operations [41,42], and bilateral filtering [38];
- ACDC is designed and developed to cope with the analysis of stacks of time-lapse microscopy images in real-time;
- ACDC does not require any training phase, and represents a reliable solution even without the availability of large-scale annotated datasets.

The manuscript is structured as follows. Section 2 describes the analyzed fluorescence imaging datasets, as well as the proposed method. Section 3 presents the results achieved by ACDC. Discussions and final remarks are provided in Section 4.

## 2. Materials and Methods

In this section we first present the fluorescence imaging datasets analyzed in this work, then we describe the pipeline at the basis of ACDC.

### 2.1. Fluorescence Microscopy Imaging Data

#### 2.1.1. Vanderbilt University Dataset

This dataset (VU) collected time-lapse microscopy images from two experiments performed in the Vanderbilt High-Throughput Screening (HTS) Core Facility at Vanderbilt University (Nashville, TN) with assistance provided by Dr. Joshua A. Bauer. All images were acquired by using Molecular Devices (San Jose, CA, USA) from ImageXpress Micro XL using a PCO.EDGE 5.5 CMOS camera with a 2560 × 2160 image sensor format (6.5 × 6.5 micron pixel size). The center 2160 × 2160 pixels were extracted during plate acquisition to ensure adequate illumination uniformity. Images were obtained using a 10× objective in the red channel (Cy3) and sometimes in the green channel (FITC) for the same well and location (overlapping information). Pixel intensity from the camera has a range of 12 bits stored in a 16-bit format. The PC-9 human lung adenocarcinoma cell line used in these studies had previously been engineered to express histone 2B conjugated to monomeric red fluorescent protein and geminin 1–110 fused to monomeric azami green [43–45]. The total number of the analyzed images with the corresponding manual cell segmentation was 46, related to two distinct experiments. The manual annotation was performed with a custom MatLab tool by a biologist (R.B.) and then validated by another expert with expertise in biochemistry and cancer biology (D.R.T.). For images with sparse cell nuclei, the manual procedure took approximately 1-2 minutes, while it took 30-40 minutes on images with high cell coverage. Two examples of input images are shown in Figure 1.

**Figure 1.**
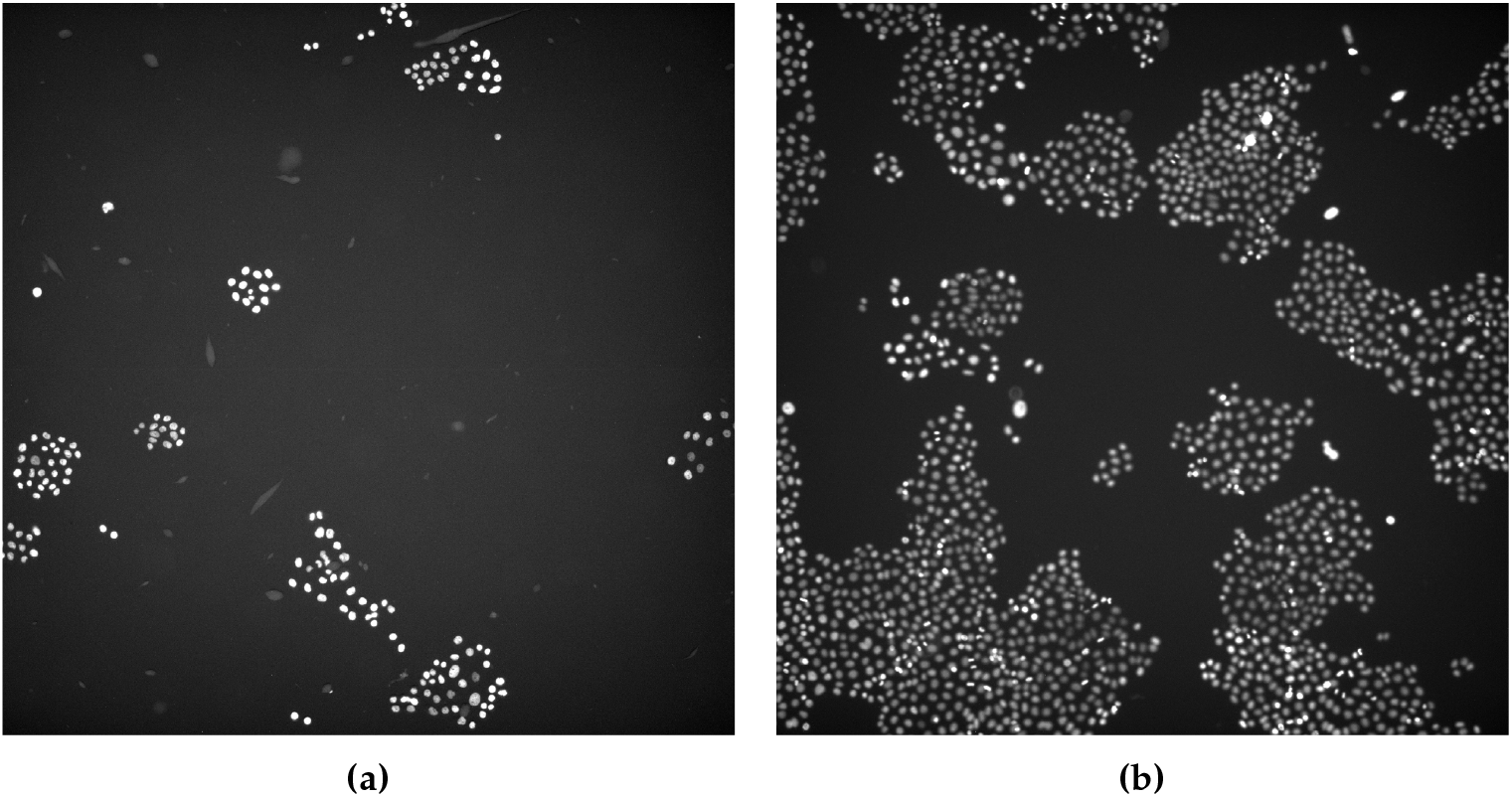
Examples of the analyzed microscopy fluorescence images provided by the Department of Biochemistry of the VU. The images were displayed by automatically adjusting the brightness and contrast according to an histogram-based procedure.

#### 2.1.2. 2018 Data Science Bowl

With the goal of validating ACDC on imaging data coming from different sources, we considered a selection of the training set of the Data Science Bowl (DSB) dataset [37,46], which was a competition organized by Kaggle (San Francisco, CA, USA). We used only the human-annotated training set because the gold standard for the test set is not publicly provided by the organizers. The annotations were manually performed by a team of expert biologists in a collaborative manner, where a single expert outlined the nuclei and the other collaborators reviewed the result.

The goal of the DSB regarded the detection of the nuclei of the cells to identify each individual cell in a sample, a mandatory operation to understand the underlying biological processes. This dataset includes a large number of labelled nuclei images acquired under a variety of conditions, magnification, and imaging modality (i.e., bright-field and fluorescence microscopy). Image size varies among 256 × 256, 256 × 320, 260 × 347, 360 × 360, 512 × 640, 520 × 696, 603 × 1272, and 1040 × 1388 pixels.

The DSB dataset aimed at evaluating the generalization capabilities of computational methods when dealing with significantly different data. Therefore, to run tests with ACDC, we first extracted the fluorescence microscopy images from the training set; then, we selected images where the maximum size of the region cells segmented in the ground truth was equal or less than 1000 pixels to obtain a magnification factor roughly comparable to the Vanderbilt University dataset, for a total of 301 images. According to the work presented in [37], we considered only the small fluorescent nuclei images (e.g., those obtained using a microscope objective of 10 × or 20 ×), which are very common in biomedical research.

Three examples of input images with highly different characteristics are shown in Figure 2.

**Figure 2.**
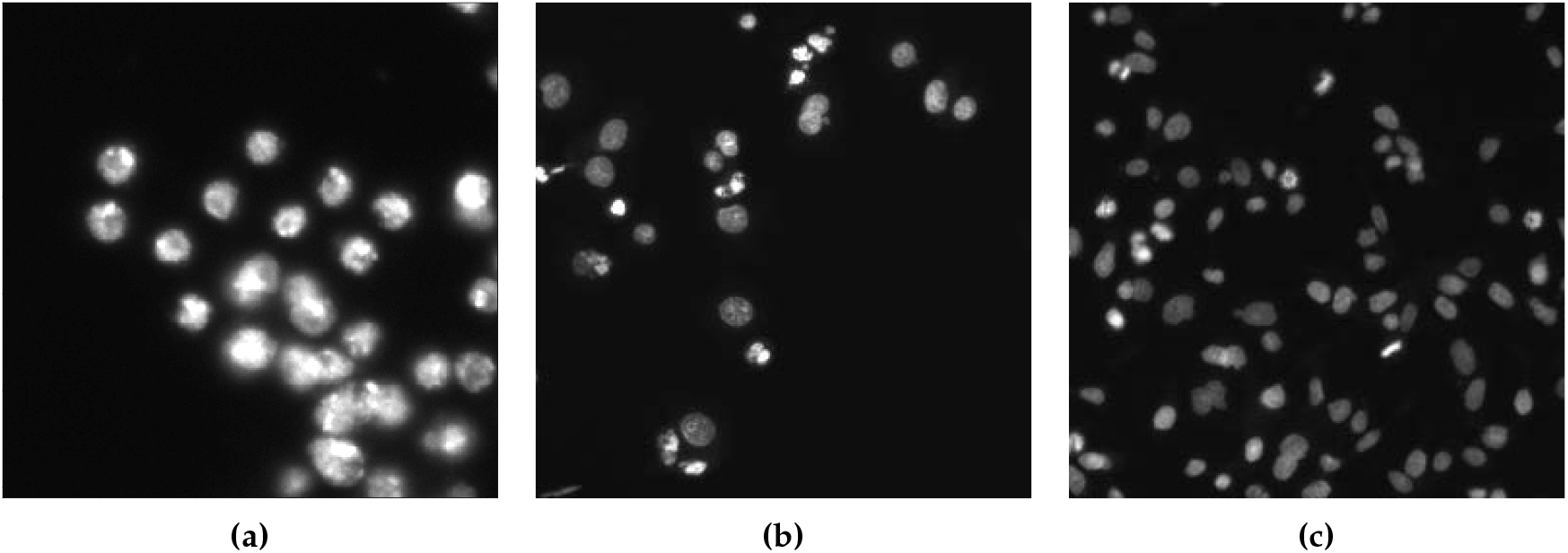
Examples of the analyzed microscopy fluorescence from the DSB dataset.

Finally, Figure 3 shows the boxplots the reveal the high variability of the analyzed microscopy imaging datasets, in terms of the number of cells and cell coverage, to support the validity of the experiments carried out in this study.

**Figure 3.**
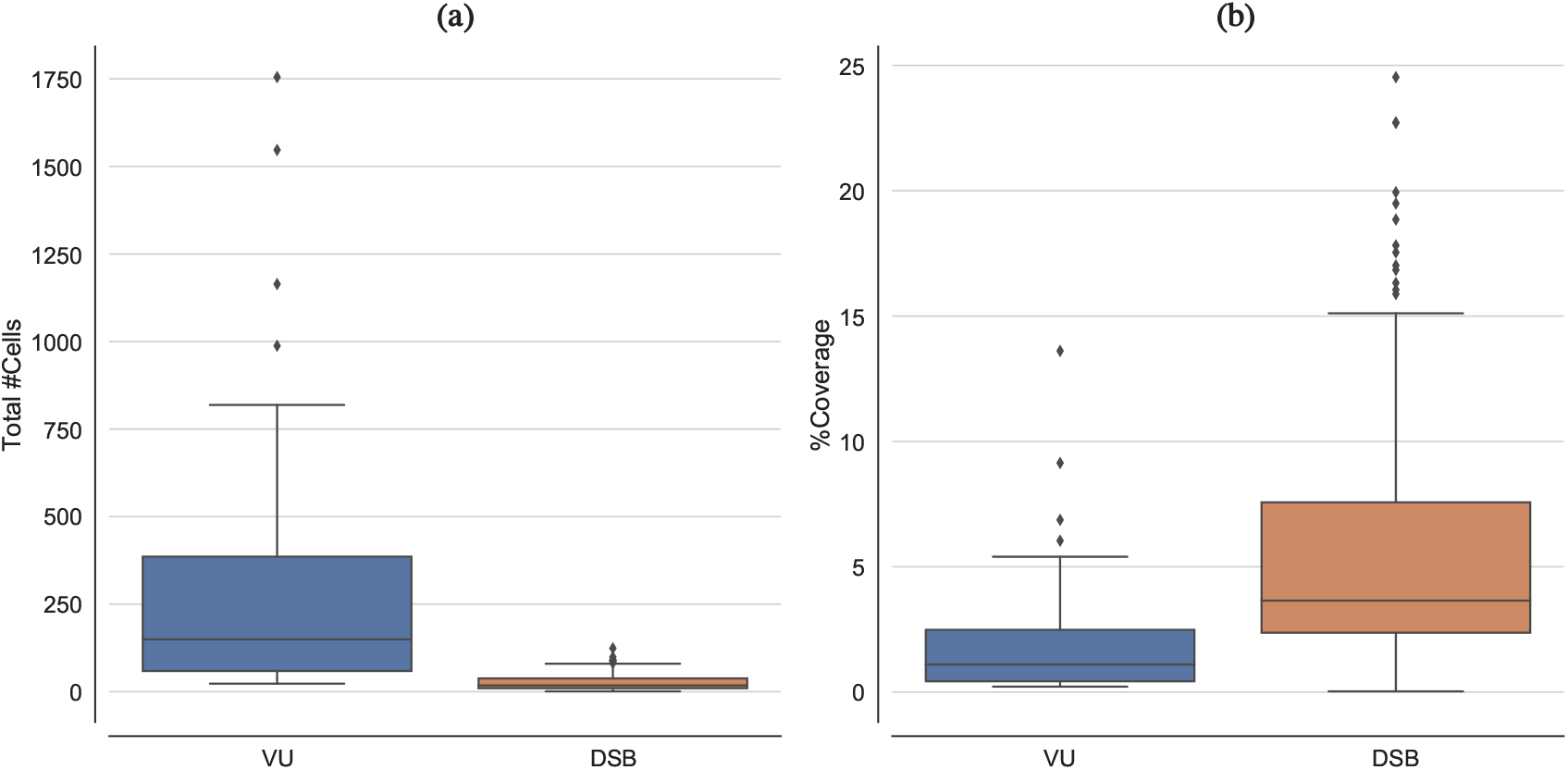
Boxplots depicting the distribution for both the analyzed datasets in terms of: **(a)** total number of cells, and **(b)** coverage of the cell nuclei regions.

### 2.2. ACDC: a Method for the Automatic Cell Detection and Counting

Advances in optics and imaging systems have enabled biologists to visualize live-cell dynamic processes by time-lapse microscopy images. However, the imaging data recorded during even a single experiment may consist of hundreds of objects over thousands of images, which makes manual inspection a tedious, time-consuming and inaccurate option. Traditional segmentation techniques proposed in the literature generally exhibit low performance on live unstained cell images. These limitations are mainly due to low contrast, intensity-variant, and non-uniform illuminated images [1]. Therefore, novel automated computational tools tailored to quantitative system-level biology are required.

ACDC is a method designed for time-lapse microscopy that aims at overcoming the main limitations of the literature methods [10,19,47], especially in terms of efficiency and execution time, by means of a fully automatic strategy that allows for reproducible measurements [48]. Each step of the pipeline underlying ACDC has been carefully optimized to reduce the running time required by large size images. We also provide two distributed versions of ACDC that allow the users to analyse a stack of images in real-time.

The processing pipeline of ACDC exploits classic image processing techniques in a smart fashion, enabling feasible analyses in real laboratory environments. Figure 4 outlines the overall flow diagram of the ACDC segmentation pipeline, as described hereafter. It is worth noting that the number of parameters that need to be set in the underlying image processing operations involves only the kernel size of spatial filters and the structuring element sizes in morphological operations, providing a reliable yet simple solution. Nevertheless, these parameters allow the user to have control over the achieved cell segmentation results. Unlike DCNN-based black- or opaque-boxes, ACDC offers an interpretable model for biologists that may conveniently adjust the parameters values according to the cell lines under investigations. Differently from supervised Machine Learning approaches [11,18,22], ACDC does not require any training phase, thus representing a reliable and practical solution even without the availability of large-scale annotated datasets.

**Figure 4.**
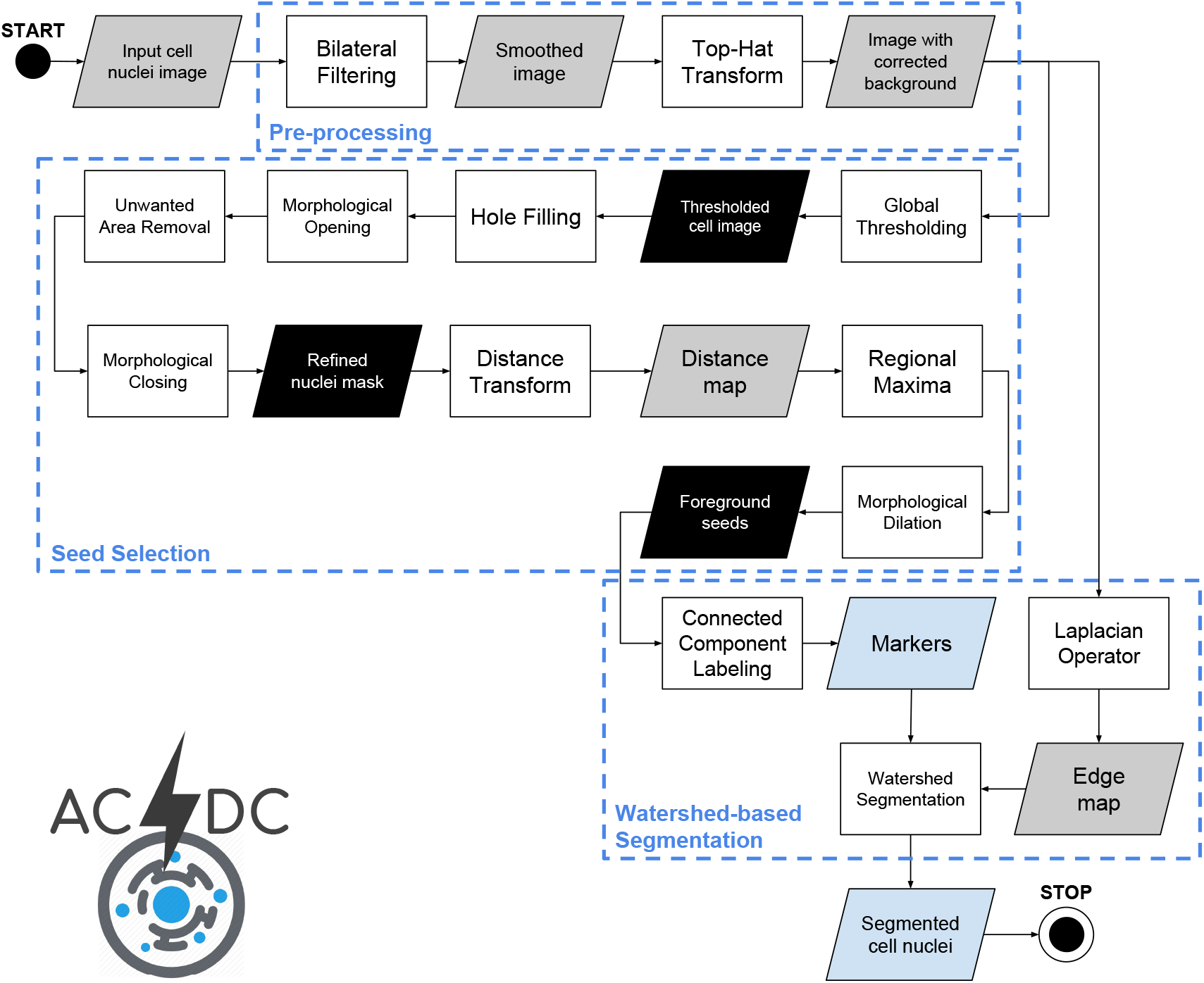
Flow diagram of the ACDC pipeline. The gray, black and light-blue data blocks denote gray-scale images, binary masks and information extracted from the images, respectively. The three macro-blocks represent the three main processing phases, namely: pre-processing, seed selection, and watershed-based segmentation.

#### 2.2.1. Pre-processing

The input microscopy image is pre-processed to yield a convenient input to the downstream watershed-based segmentation by means of the following steps:

1. Application of bilateral filtering that allows for denoising the image I while preserving the edges by means of a non-linear combination of nearby image values [38]. This noise-reducing smoothing filter combines gray levels (colors) according to both a geometric closeness function *c* and a radiometric (photometric) similarity function s. This combination is used to strengthen near values with respect to distant values in both spatial and intensity domains. This simple yet effective strategy allows for contrast enhancement [49]. Bilateral filter has been shown to work properly in fluorescence imaging even preserving the directional information, such as in the case of the F-actin filaments [50]. This denoising technique was effectively applied to biological electron microscopy [51], as well as to cell detection [52], revealing better performance—compared to low-pass filtering—in noise reduction without removing the structural features conveyed by strong edges. The most commonly used version of bilateral filtering is the shift-invariant Gaussian filtering, wherein both the closeness function *c* and the similarity function *s* are Gaussian functions of the Euclidean distance between their arguments [38]. With more details, *c* is radially symmetric: 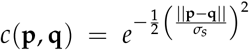. Consistently, the similarity function *s* can be defined as: 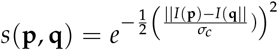. In ACDC we set *σ_c_* = 1 and *σ_s_* = *σ*_global_ (where *σ*_global_ is the the standard deviation of the input image 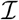) for the standard deviation of the Gaussian functions *c* and *s*, respectively. This smart denoising approach allows us to keep the edge sharpness while reducing the noise of the processed image, so avoiding cell region under-estimation.
2. Application of top-hat transform for background correction with a binary circular structuring element (radius: 21 pixels) on the smoothed image. This operation accounts for non-uniform illumination artifacts, by extracting the nuclei from the background. The white top-hat transform is the difference between the input image *I* and the opening of 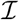 with a gray-scale structuring element 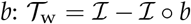 [53].

The results of the pre-processing images applied to Figures 1a and 2a are shown in Figures 5a and 5b, respectively. For the pre-processing step, ACDC requires only 3 parameters, namely: *σ_c_* and *σ_s_* for the bilateral filtering, and a structuring element for the hat-top transform.

**Figure 5.**
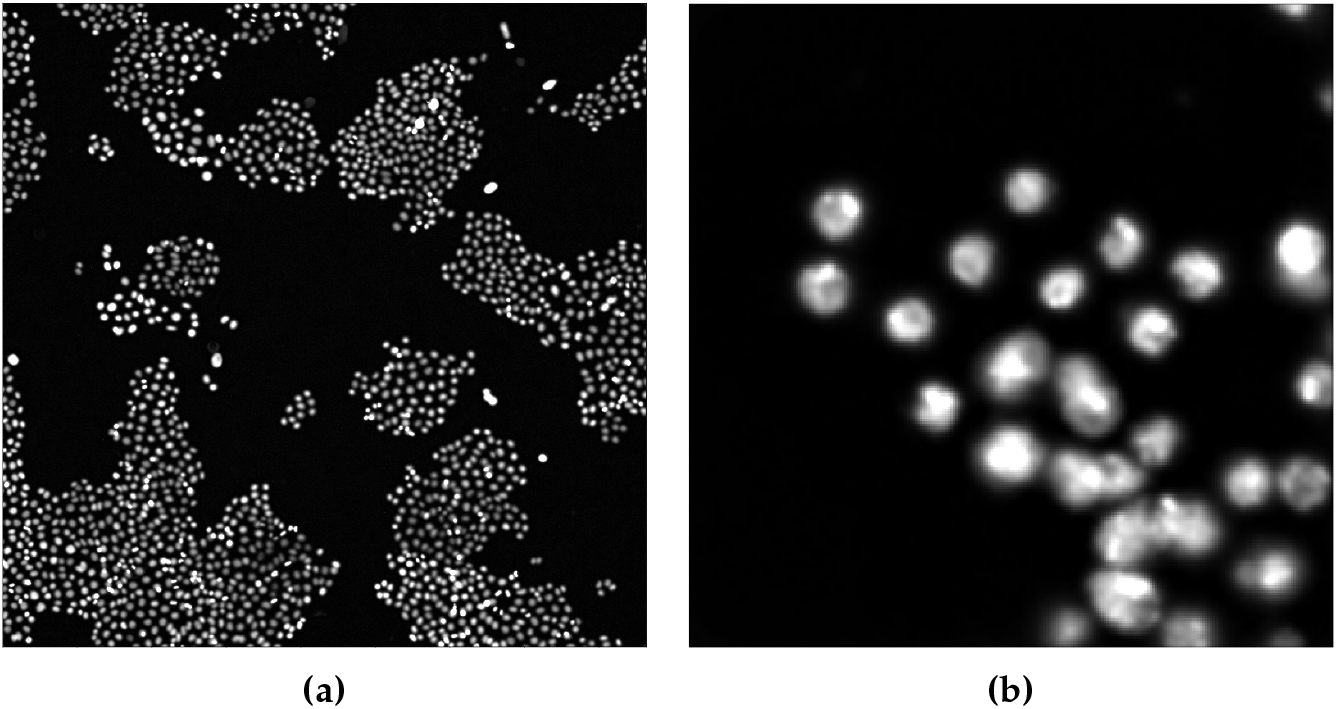
Result of the application of the pre-processing steps on the images shown in Figure 1a **(a)** and Figure 2b **(b)**.

#### 2.2.2. Nucleus Seed Selection

The following steps are executed to obtain a reliable seed selection, so that the cells nuclei can be accurately extracted from the pre-processed images:

1. A thresholding technique has to be first applied to detect the cell regions. Both global and local thresholding techniques aim at separating foreground objects of interest from the background in an image, considering differences in pixel intensities [54]. Global thresholding determines a single threshold for all pixels and works well if the histogram of the input image contains well-separated peaks corresponding to the desired foreground objects and background [55]. Local adaptive thresholding techniques estimate the threshold locally over sub-regions of the entire image, by considering only a user-defined window with a specific size and exploiting local image properties to calculate a variable threshold [53,54]. These algorithms find the threshold by locally examining the intensity values of the neighborhood of each pixel according to image intensity statistics. To avoid unwanted pixels in the thresholded image, mainly due to small noisy hyper-intense regions caused by non-uniform illumination, we apply the Otsu global thresholding method [55] instead of local adaptive thresholding based on the mean value in a neighborhood [56]. Moreover, global threshold techniques are significantly faster than local adaptive strategies.
2. Hole filling is applied to remove possible holes in the detected nuclei due to small hypo-intense regions included in the nuclei regions.
3. Morphological opening (using a disk with 1-pixel radius as a structuring element) is used to remove loosely connected-components, such as in the case of almost overlapping cells.
4. Unwanted areas are removed according to the connected-components size. In particular, the detected candidate regions with areas smaller than 40 pixels are removed to refine the achieved segmentation results by robustly avoiding false positives.
5. Morphological closing (using a 2-pixel radius circular structuring element) is applied to smooth the boundaries of the detected nuclei and avoid the under-estimation of the detected nuclei regions.
6. The approximate Euclidean distance transform (EDT) from the binary mask, achieved by applying the Otsu algorithm and refined by using the previous 3 steps, is used to obtain the matrix of distances of each pixel to the background by exploiting the *ℓ*_2_ Euclidean distance [57] (with a 5 × 5 pixel mask for a more accurate distance estimation). This algorithm calculates the distance to the closest background pixel for each pixel of the source image. Let 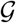 be a regular grid and 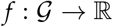 an arbitrary function on the grid, called a sampled function [58]. We define the distance transform 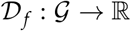 of *f* as:

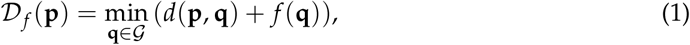

where *d*(**p, q**) is a measure of the distance between the pixels **p** and **q**. Owing to the fact that cells have a pseudo-circular shape, we used the Euclidean distance, achieving the EDT of *f*. In the case of binary images, with a set of points 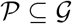, the distance transform 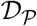 is a real-valued image of the same size:

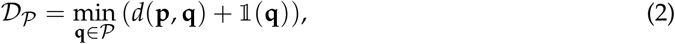

where:

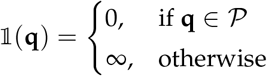

is an indicator function for the membership in 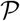 [58]. The computed distance map is normalized by applying contrast linear stretching to the full 8-bit dynamic range.
7. Regional maxima computation allows for estimating foreground peaks on the normalized distance map. Regional maxima are connected-components of pixels with a constant intensity value, whose external boundary pixels have all a lower intensity value [42]. The resulting binary mask contains pixels that are set to 1 for identifying regional maxima, while all other pixels are set to 0. A 5 × 5 pixel square was employed as structuring element.
8. Morphological dilation (using a 3-pixel radius circular structuring element) is applied to the foreground peaks previously detected for better defining the foreground regions and merging neighboring local minima into a single seed point. The segmentation results on Figures 5a and 5b are shown in Figures 6a and 6b, respectively. The detail in Figure 5a shows that ACDC is highly specific to cell nuclei detection, discarding non-cell regions related to acquisition artifacts.

**Figure 6.**
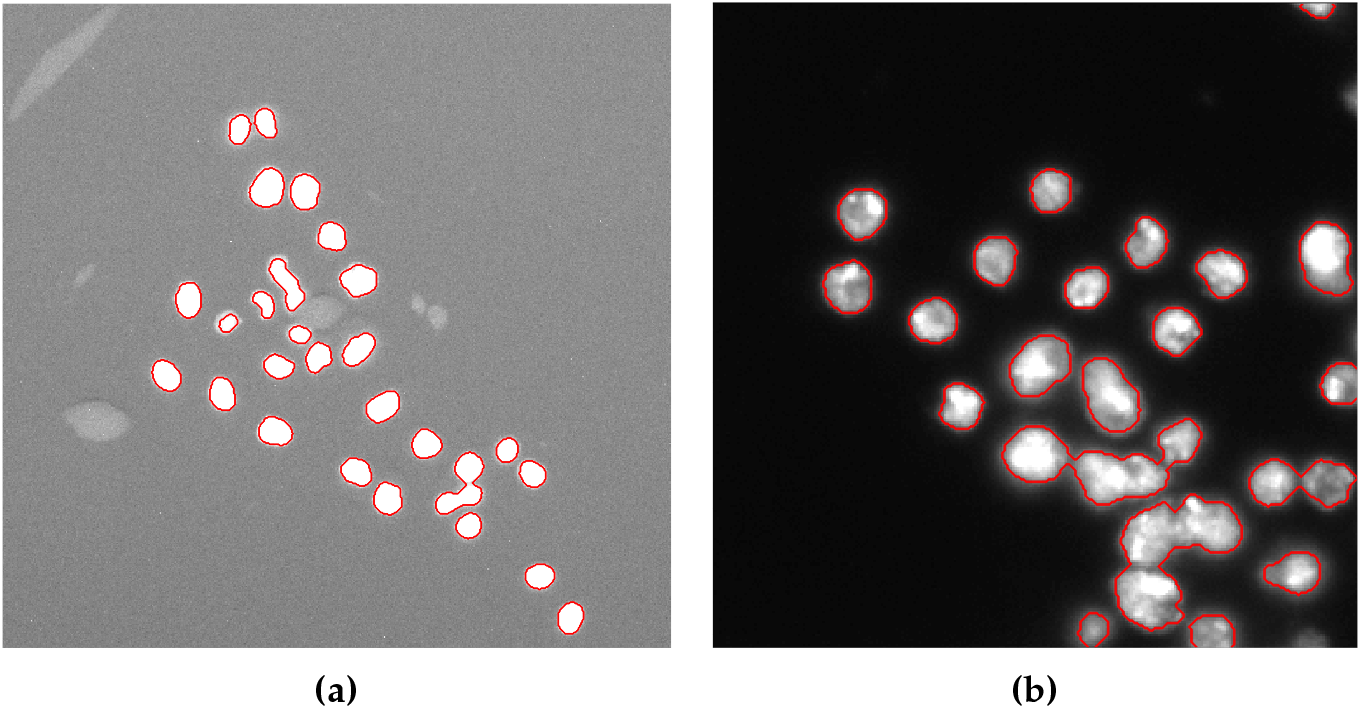
Segmented images obtained after the refinement steps applied to the image shown in Figure 5a **(a)**, and to a sub-image of Figure 5b **(b)**.

For the Nucleus Seed Selection step of the pipeline, the user can set the following parameters: a structuring element for the morphological opening; a structuring element for the morphological closing; a structuring element for the morphological dilation; the minimum size (in pixels) for the unwanted area removal.

#### 2.2.3. Cell Nuclei Segmentation Using the Watershed Transform

The watershed transform [39] is one of the most used approaches in cell image segmentation [7], while it was originally proposed in the field of mathematical morphology [42].

The intuitive description of this transform is straightforward: assuming an image as a topographic relief, where the height of each point is directly related to its gray level, and considering rain gradually falling on the terrain, then the watersheds are the lines that separate the resulting catchment basins [40]. This technique is valuable because the watershed lines generally correspond to the most significant edges among the markers [41] and are useful to separate overlapping objects, such as in the case of the nuclei separation in cell segmentation in human-derived cardiospheres (i.e., 3D clusters of cardiac progenitor cells) [59]. Even when no strong edges between the markers exist, the watershed method is able to detect a contour in the area. This contour is detected on the pixels with higher contrast [39]. As a matter of fact, edge-based segmentation techniques—which strongly rely on local discontinuities in gray levels—often do not yield unbroken edge curves, thus heavily limiting their performance in cell segmentation [12]. Unfortunately, it is also well-known that the watershed transform may be affected by over-segmentation issues, thus requiring further processing [60].

From a computational perspective, the watershed algorithm analyzes a gray-scale image by means of a flooding-based procedure. Since the flooding process is performed on either a gradient image or edge map, the basins should emerge along the edges. As a matter of fact, during the watershed process, the edge information allows for a better discrimination of the boundary pixels with respect to the original image. Finally, only the markers of the resulting foreground cells are selected. ACDC uses an efficient version of the watershed algorithm that exploits a priority queue to store the pixels according to the pixel value (i.e., the height in the gray-scale image landscape) and the entry order into the queue (giving precedence to the closest marker). More specifically, during the flooding procedure, this process sorts the pixels in increasing order of their intensity value by relying on a breadth-first scan of the plateaus based on a first-in-first-out data structure [40].

Although the watershed transform can detect also weak edges, it may not accurately detect the edge of interest in the case of blurred boundaries [60]. This sensitivity to noise could be worsened by the use of high pass filters to estimate the gradient and the edges, which amplify the noise. We address this issue by formerly applying the bilateral filter that reduces the halo effects [38]. Accordingly, we implemented the following steps:

1. Connected-component labeling [61] of the foreground region binary mask for encoding the markers employed in the following watershed algorithm.
2. Laplacian operator for producing the edge image [53] to feed the edge map as input to the watershed transform.
3. Watershed segmentation on the edge image according to the previously defined markers [62,63]. ACDC does not require any settings for the Cell Nuclei Segmentation Using the Watershed Transform step.

#### 2.2.4. Implementation Details

The sequential version of ACDC has been entirely developed using the Python programming language (version 2.7.12), exploiting the following libraries and packages: NumPy, SciPy, OpenCV, scikit-image [64], and Mahotas [65]. The resulting processing pipeline makes use of classic image processing techniques in a smart fashion [66], thus enabling an efficient and feasible solution in time-lapse microscopy environments.

For laboratory feasibility purposes, an asynchronous job queue, based on a distributed message passing paradigm—namely Advanced Message Queuing Protocol (AMQP)—was developed using Celery [67] (implementing workers that execute tasks in parallel) and RabbitMQ [68] (exploited as a message broker to handle communications among workers) for leveraging modern multi-core processors and computer clusters.

We also developed a Parent-Workers strategy using mpi4py, which provides bindings of the Message Passing Interface (MPI) specifications for Python to leverage multi-core and many-core resources [69]. The distributed strategy used to accelerate ACDC is similar to that employed in [70–72], where the Parent allocates the resources and orchestrates the workers, which run ACDC to analyze the assigned images. This distributed version of ACDC is 3.7× faster than the sequential version by exploiting 6 cores of a CPU Intel Core E5-2650 v4 (clock 2.2 GHz).

### 2.3. Segmentation Evaluation Metrics

The accuracy of the achieved segmentation results 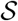 was quantitatively evaluated with respect to the real measurement—i.e., the ground truth 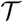 obtained manually by an experienced biologist—by using the Dice Similarity Coefficient (DSC):

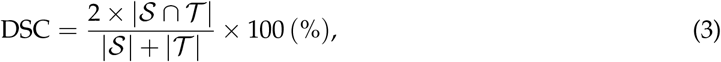

as well as the Intersection over Union (IoU) metrics, also known as Jaccard coefficient:

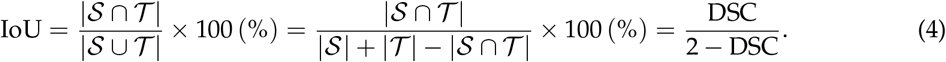

## 3. Results

### 3.1. ACDC performance

In this section we present the results obtained with ACDC on the VU dataset and the 2018 DSB training dataset [37,46]. Figure 7 shows an example of results obtained on VU and DSB datasets, where the detected cells are displayed with different colors to highlight the separation among overlapping and merging cells. We note that the analyzed images are characterized by a considerable variability in terms of cell density. Figure 8 shows a representative case for both the VU and DSB datasets, where the results of ACDC are slightly different from the gold standard. In the case of the VU dataset (Figure 8a), it is clear that some groups of small cells, characterized by very similar intensity values without strong discontinuities are detected by ACDC as a single connected-component (orange arrows). On the contrary, for the 2018 DSB dataset (Figure 8b), ACDC is capable of accurately separating groups of cells that were erroneously delineated as single connected-components in the gold standard (green arrows); moreover, the spurious speckles included in the 2018 DSB gold standard (highlighted by blue dashed boxes), are not detected by ACDC since these very small regions are characterized by hypo-intense fluorescence levels.

**Figure 7.**
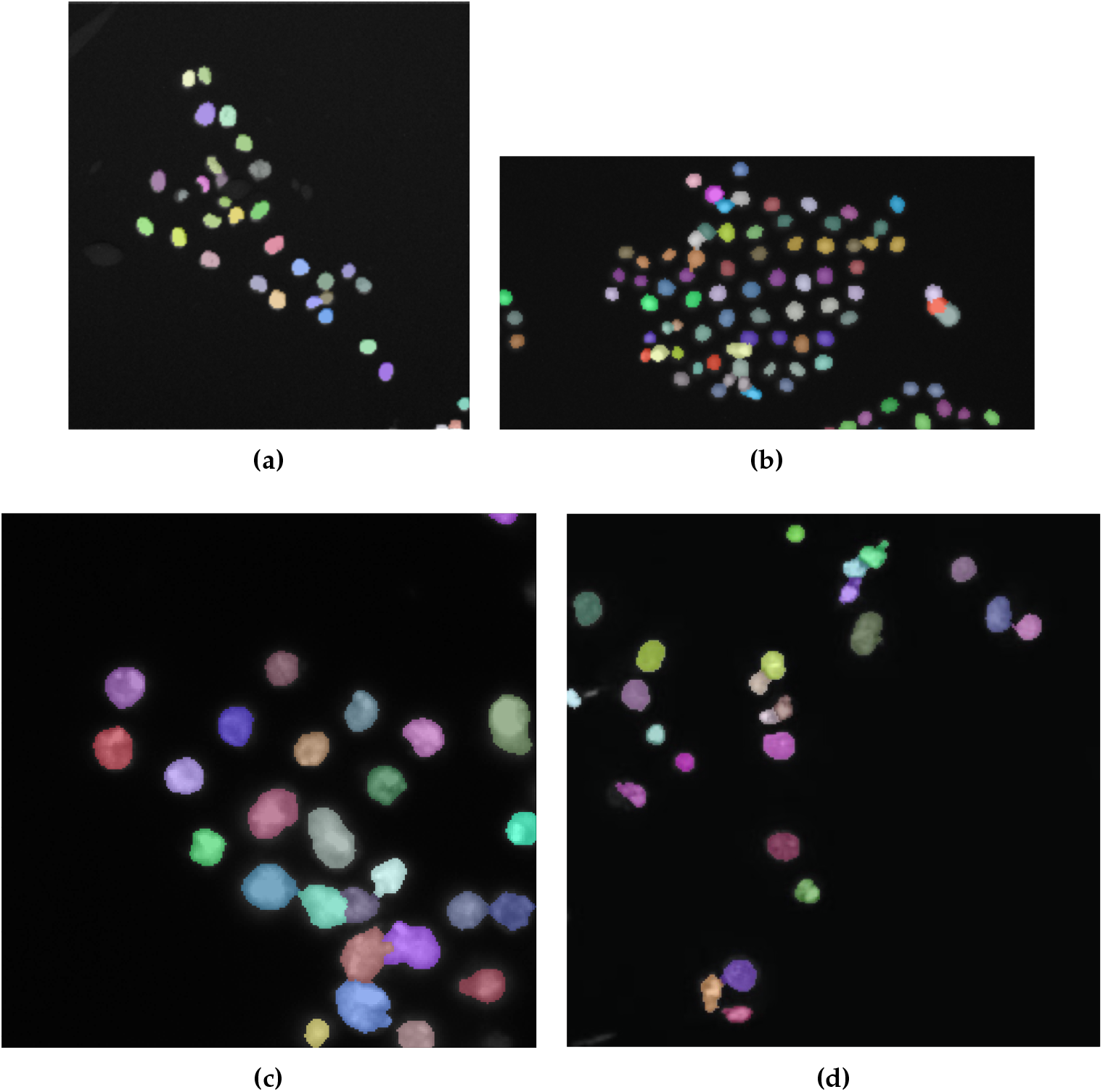
Examples of cell nuclei segmented by ACDC considering sub-images of Figures 1a and 1b **(a)-(b)**, and the whole images in Figures 2a and 2b **(c)-(d)**. The cell nuclei images were over-imposed onto the original fluorescence images with alpha-blending (*α* = 0.4)

**Figure 8.**
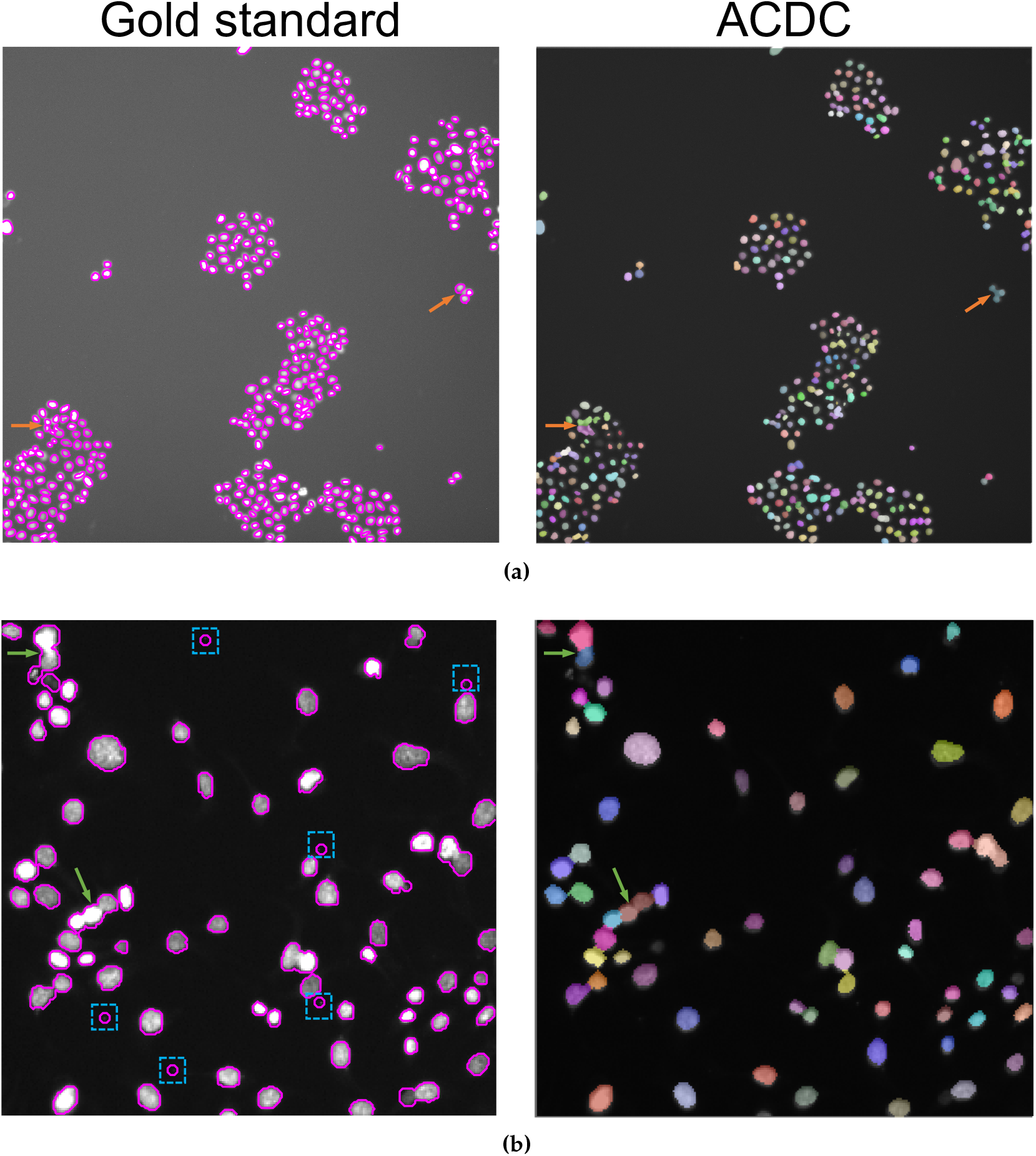
Comparison of the gold standard cell nuclei segmentation (magenta contour in the left images) against the automated result obtained by ACDC (segmented nuclei over-imposed onto the original fluorescence images with alpha-blending in the right images): **(a)** VU dataset, where the orange arrows denote errors in the split of clustered cell nuclei; **(b)** DSB dataset, where the green arrows denote groups of cells that were erroneously delineated as unique connected-components in the gold standard and the blue dashed boxes represent spurious speckles that are not detected by ACDC.

The assessment of cell count was quantified by means of the Pearson coefficient to measure the linear correlation between the automated and manual cell counts. The accuracy of the achieved segmentation results was evaluated by using the DSC and the IoU metrics. Table 1 reports the results—by also comparing the performance of ACDC with and without bilateral filtering—achieved on the VU and 2018 DSB datasets, which are supported by the scatter plots in Figure 9 that reports the results concerning the case of the bilateral filter. The achieved high *ρ* coefficient values confirm the effectiveness of the proposed approach, according to a validation performed against the manual cell counting, which is considered as the gold standard.

**Figure 9.**
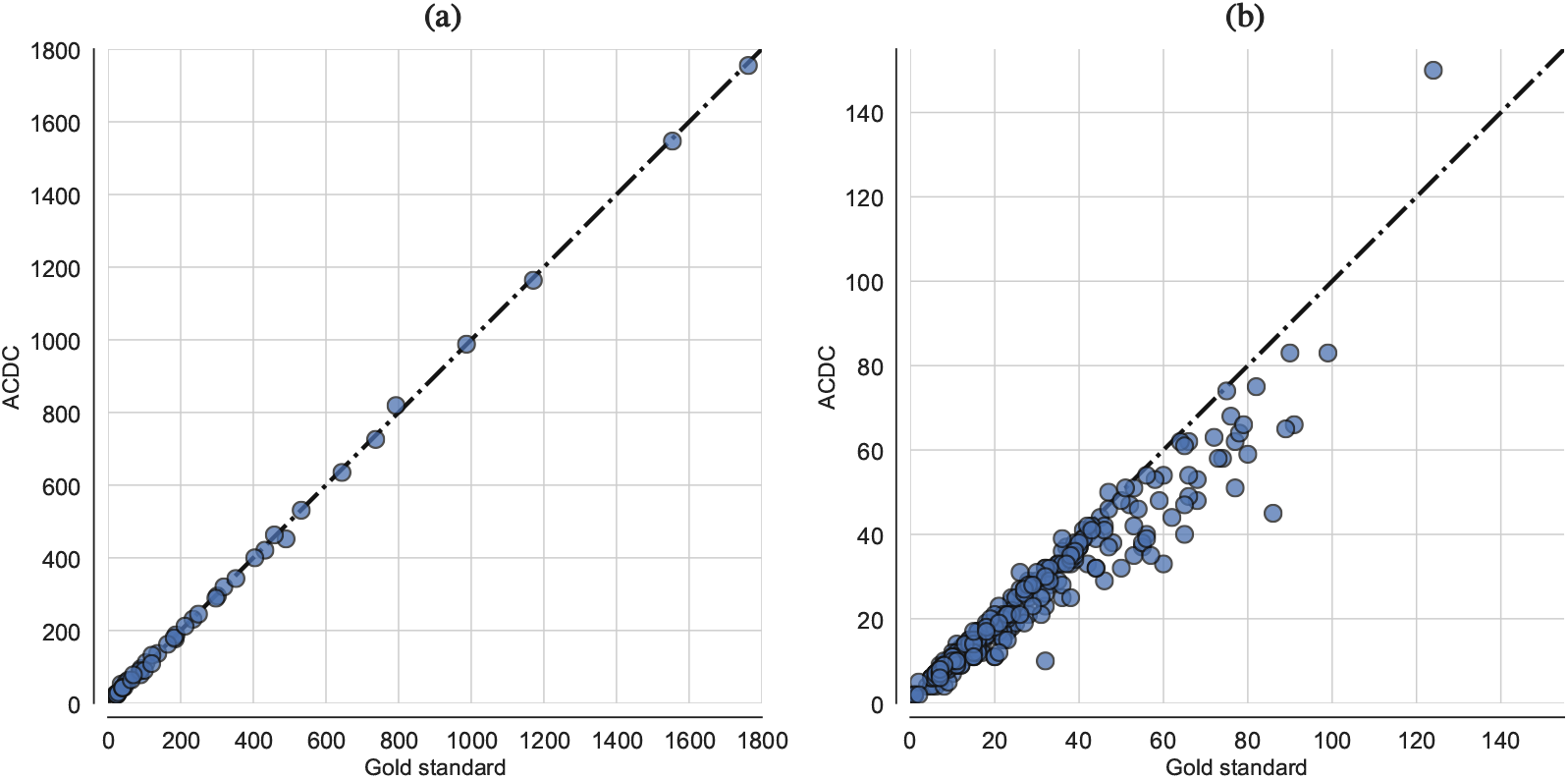
Scatter plots depicting ACDC results compared to the gold standard in terms of cell nuclei counting in the case of: **(a)** time-lapse fluorescence images from the VU dataset; **(b)** small fluorescent nuclei images from the DSB dataset. The equality line through the origin is drawn as a dashed line.

**Table 1.** Evaluation metrics on cell counting and segmentation achieved by ACDC (with and without bilateral filtering) on the analyzed time-lapse microscopy VU and 2018 DSB datasets, comprising 46 and 301 images, respectively. The results for the DSC and IoU metrics, as well as the execution time measurements, are expressed as mean value ± standard deviation.

| Method | Dataset | Pearson coeff. ($p$-value) | DSC (%) | IoU (%) | Exec. time (s) |
| --- | --- | --- | --- | --- | --- |
| ACDC (without bilateral filter) | VU | $\rho = 0.99$ ($p = 2.5 \times 10^{-74}$) | $75.86 \pm 5.98$ | $61.45 \pm 7.47$ | $3.98 \pm 0.10$ |
| ACDC (with bilateral filter) | VU | $\rho = 0.99$ ($p = 6.6 \times 10^{-74}$) | $76.84 \pm 6.71$ | $62.84 \pm 8.58$ | $7.49 \pm 0.30$ |
| ACDC (without bilateral filter) | DSB | $\rho = 0.96$ ($p = 1.1 \times 10^{-175}$) | $87.34 \pm 6.89$ | $77.97 \pm 9.49$ | $0.07 \pm 0.05$ |
| ACDC (with bilateral filter) | DSB | $\rho = 0.96$ ($p = 2.6 \times 10^{-169}$) | $88.64 \pm 7.41$ | $80.37 \pm 10.58$ | $0.12 \pm 0.09$ |

A strong positive correlation between the cell counts computed automatically by ACDC and the manual measurements was observed for both datasets analyzed in this study. In particular, in the case of the VU dataset, the automated cell counting is strongly consistent with the corresponding manual measurements, by denoting also a unity slope, as shown in Figure 9a. In the case of the DSB dataset (Figure 9b), the identity straight-line reveals a negative offset in the ACDC measurements. This finding means that the cell counts achieved by ACDC slightly under-estimated the gold standard in approximately 55% of cases. Besides the high variability of the fluorescence images included in the DSB dataset, this systematic error often depends on the gold standard binary masks, where the cell nuclei are not always precisely separated (since the DSB challenge was focused on the segmentation task), and consider also partial connected-components of cell nuclei smaller than 40 pixels located at the cropped image borders. Notice that no ACDC’s setting was modified, so some very small partial cell components were removed according to the morphological refinements based on the connected-component size.

The agreement between ACDC and the manual cell counting measurements can be graphically represented also by using a Bland–Altman plot [73], as reported in Figure 10 for both datasets. The Bland–Altman analysis allows for a better data visualization by showing the pairwise difference between the measurements obtained with the two methods, against their mean value. In this way, we can assess any possible relationship between the estimation errors and easily detect any outlier (i.e., observations outside the 95% limits of agreement). Figure 10 reveals that ACDC achieved reproducible results also by considering the whole range of the cell counts concerning the analyzed cell microscopy images; only about the 4% and 6% of outliers are observed in the ACDC results achieved on the VU and DSB datasets, respectively. This confirms the observations on the scatter plots in Figure 9; neither systematic discrepancy between the measurements nor bias can be observed in the VU dataset, while a negative bias is visible in the case of the DSB dataset.

**Figure 10.**
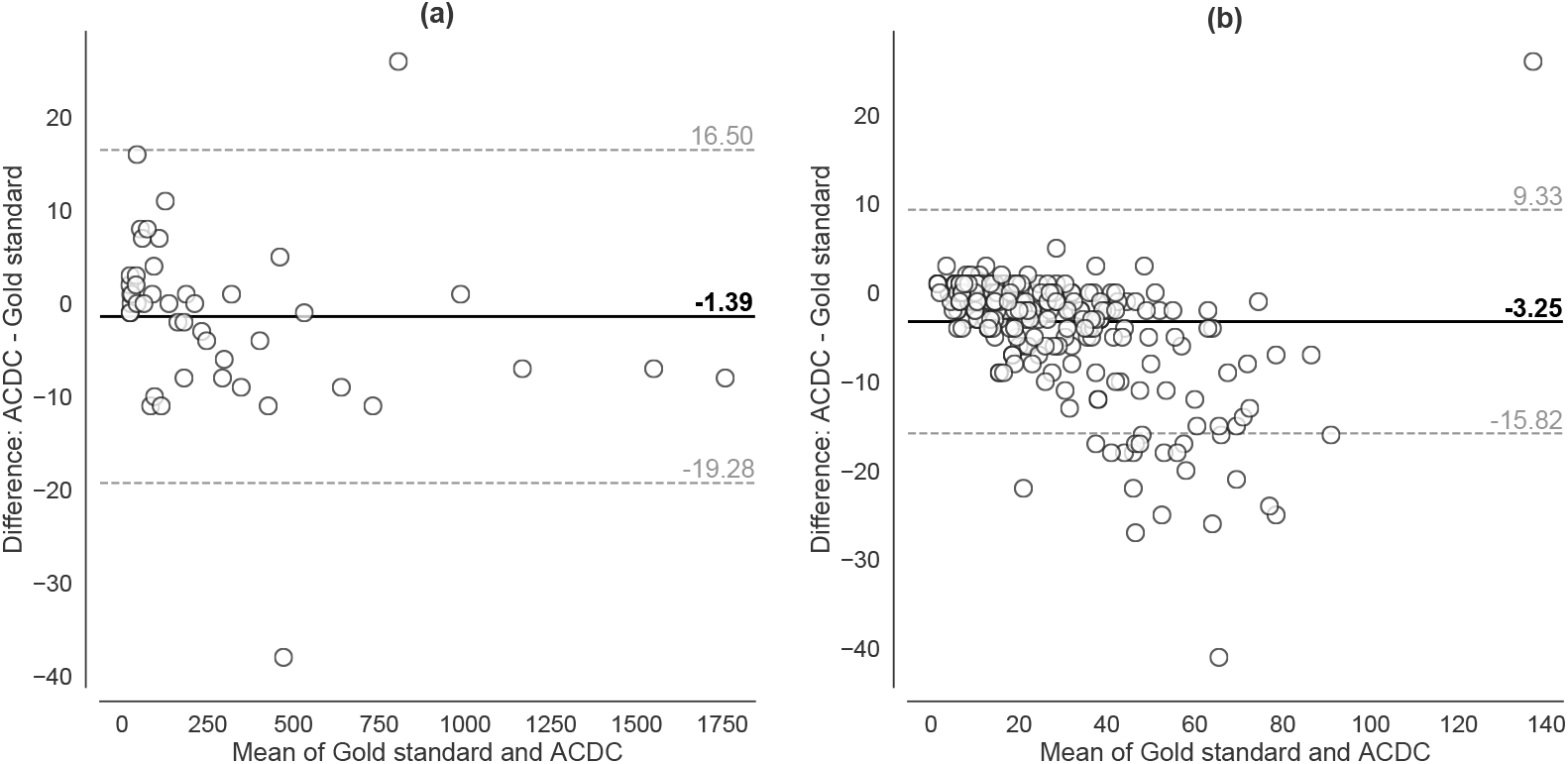
Bland–Altman plots of the cell counting measurements achieved by ACDC versus the gold standard for the **(a)** VU and **(b)** DSB datasets. Solid horizontal and dashed lines denote the mean and ±1.96 standard deviation values, respectively.

The analysis of the DSC and IoU mean values reported in Table 1, calculated on the VU images, reveals a good agreement between the segmentation results achieved by ACDC and the gold standard. The standard deviation values confirm the high variability encountered in the input datasets. In particular, in the case of the VU images, this evidence strongly depends on the density of the cells represented in the input image. As a matter of fact, the DSC and IoU metrics is highly affected by the size of the foreground regions with respect to the background. This behavior is confirmed by the results achieved on the DSB dataset, where the IoU values are considerably higher than those achieved on the VU images, even though the Pearson coefficient is slightly lower in this case. Accordingly, the high standard deviation for the DSB dataset is due to the intrinsic variety of the images—in terms of image size, zoom factor, and specimen type—included in this dataset (see Section 2.1.2). In general, the bilateral filtering allows us to achieve better segmentation performance, while no appreciable difference was found in the *ρ* values. Therefore, we incorporated the bilateral filtering into the ACDC pipeline tested in the experiments.

The mean execution times concerning the segmentation tests are shown in Table 1. These experiments were run on a personal computer equipped with a quad-core Intel Core 7700HQ (clock frequency 3.80 GHz), 16 GB RAM, and Ubuntu 16.04 LTS operating system. The computational efficiency of ACDC is confirmed in both cases, as cell detection and counting tasks can be completed in respect of the time constraints imposed by the high-throughput laboratory routine. As expected, the execution times are dependent on the image size. This trend is mainly due to the bilateral filtering operation, as it can be observed in the execution times measured on the VU dataset reported in Table 1.

### 3.2. Comparison with Other Cell Imaging Tools and Segmentation Methods

We compared the performance of ACDC against two pipelines realized with ImageJ v1.53c [8]. Since filtering operations typically represent the initial step used to enhance the subsequent segmentation, in order to assess any performance improvement introduced by low-pass filtering, we implemented the processing pipelines in ImageJ, to perform cell segmentation and counting, with and without a Gaussian filtering. To be more precise, the steps are: (*i*) Gaussian low-pass filtering; (*ii*) uneven background removal by an histogram-based rolling-ball algorithm; (*iii*) Otsu global thresholding; (iv) morphological operations including hole-filling; (v) watershed algorithm for cell nuclei separation. The pipelines have been tested on both VU and DSB datasets.

Figure 11 shows the results concerning cell counting achieved by ACDC and the two ImageJ pipelines. In the case of the VU dataset (plots a-c), the ImageJ pipelines tend to over-estimate the number of cells in the images; on the contrary, considering the DSB dataset (plots d-f), the results are comparable. The segmentation results shown in Figure 12 present a different scenario. ACDC is generally better than the ImageJ pipelines; we also note that the application of Gaussian filtering in the ImageJ pipelines does not allow us to markedly improve the segmentation outcome. However, over-segmentation can be observed when low-pass filtering is not applied.

**Figure 11.**
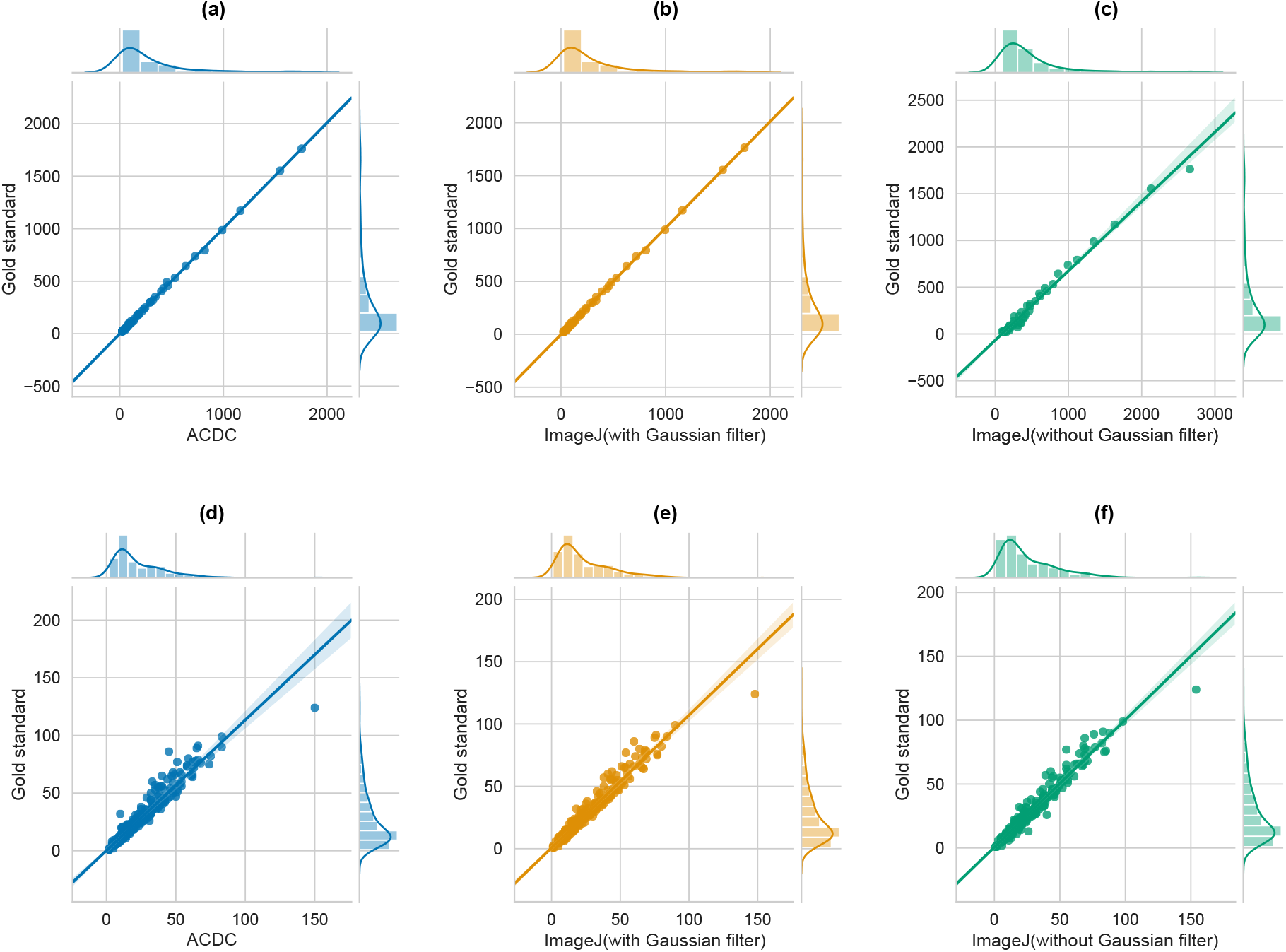
Regplots showing the scatter plots obtained by considering the number of manually detected cells (y-axis) and the number of cells automatically detected (x-axis) with ACDC (left), ImageJ with Gaussian filter (center), and ImageJ without Gaussian filter (right), along with the fitted regression model (regression line and the 95% confidence interval for that regression). Plots **(a-c)** report the results obtained on the VU dataset, while plots **(d-f)** are obtained from the DSB dataset.

**Figure 12.**
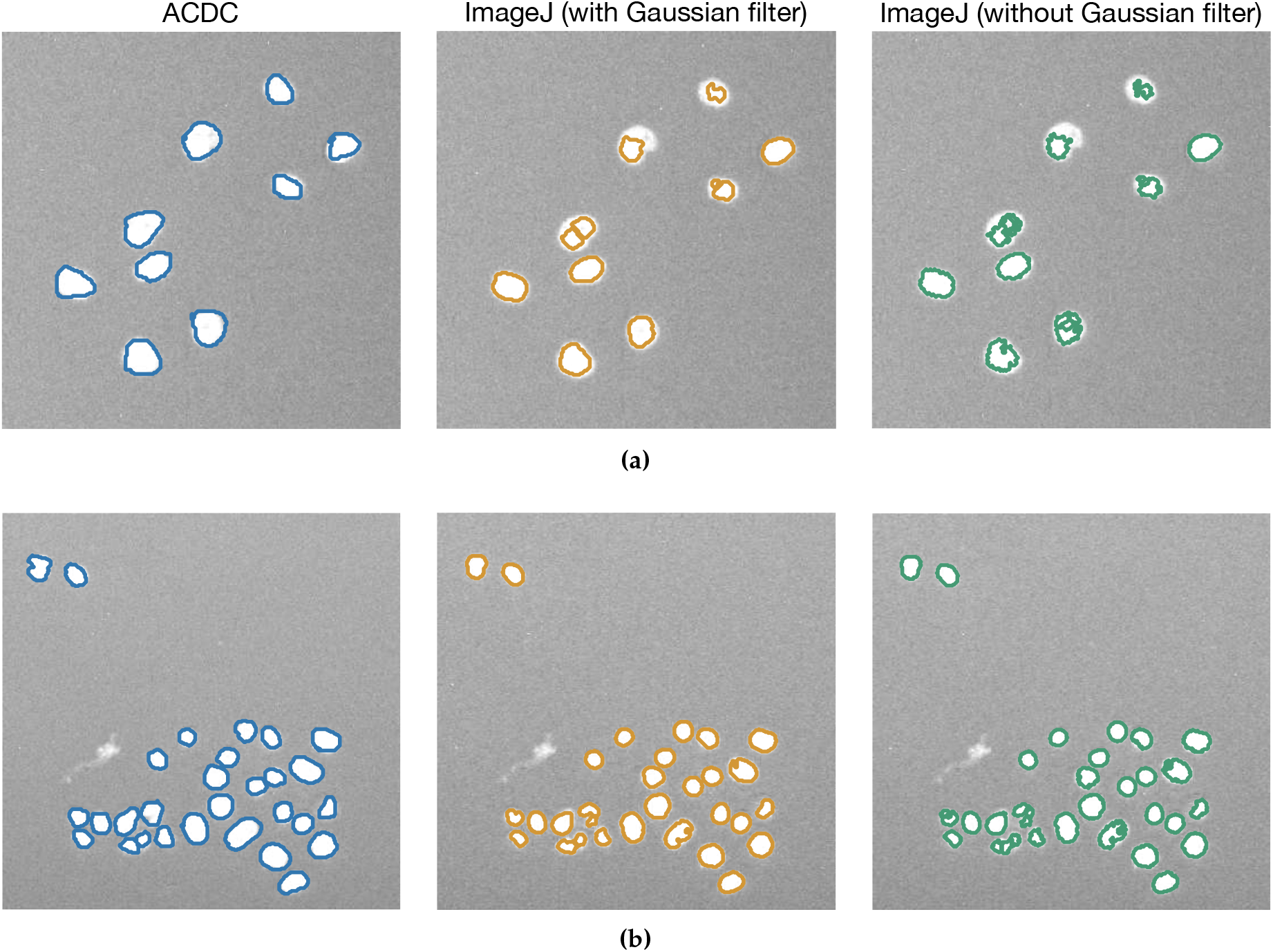
Examples of cell nuclei segmented by using ACDC and the implemented ImageJ pipelines with and without Gaussian filtering. (**a**) and (**b**) present images taken from the VU dataset, with different visual characteristics.

Table 2 reports the values of DSC and IoU achieved by ACDC and the tested ImageJ pipelines. The highest values for these metrics are obtained by ACDC both in the case of VU and DSB datasets. Moreover, the very high *ρ* coefficients indicate that all approaches are effective, when validated against the gold standard. To assess whether the difference in the achieved segmentation performance between ACDC and ImageJ with Gaussian filter is statistically significant, we performed a two-sided Wilcoxon signed rank test on both paired DSC and IoU results [74], with the null hypothesis that the samples come from continuous distributions with equal medians, and considering a significance level of 0.05. The results of the test confirmed that both the achieved DSC and IoU values are statistically different in the case of ACDC and ImageJ with Gaussian filtering (*p* < 0.001 and *p* < 0.01 for the VU and DSB datasets, respectively).

**Table 2.** Comparison of ACDC and the tested ImageJ pipelines in terms of cell nuclei counting and segmentation. The results are expressed as mean value ± standard deviation.

| Method | Dataset | Pearson coeff. ($p$-value) | DSC (%) | IoU (%) |
| --- | --- | --- | --- | --- |
| ACDC | VU | $\rho = 0.99$ ($p = 6.6 \times 10^{-74}$) | $76.84 \pm 6.71$ | $62.84 \pm 8.58$ |
| ImageJ (with Gaussian filter) | VU | $\rho = 0.99$ ($p = 9.5 \times 10^{-75}$) | $74.97 \pm 6.17$ | $60.32 \pm 7.62$ |
| ImageJ (without Gaussian filter) | VU | $\rho = 0.99$ ($p = 5.3 \times 10^{-44}$) | $74.43 \pm 6.20$ | $59.62 \pm 7.61$ |
| ACDC | DSB | $\rho = 0.96$ ($p = 2.6 \times 10^{-169}$) | $88.64 \pm 7.41$ | $80.37 \pm 10.58$ |
| ImageJ (with Gaussian filter) | DSB | $\rho = 0.97$ ($p = 5.7 \times 10^{-205}$) | $86.50 \pm 6.86$ | $76.78 \pm 9.66$ |
| ImageJ (without Gaussian filter) | DSB | $\rho = 0.97$ ($p = 3.4 \times 10^{-207}$) | $86.68 \pm 7.75$ | $77.23 \pm 10.86$ |

As a final test, we considered a recently published work that takes into account the SNP HEp-2 Cell Dataset (SNPHEp-2) [75]. SNPHEp-2 is composed of images acquired using a monochrome high dynamic range microscopy camera, equipped with a plan-Apochromat 20× /0.8 objective lens and an LED illumination source, resulting in a dataset considerably different compared to VU and DSB datasets. Specifically, we compared ACDC against CellProfiler [10], Marker-Controlled Watershed algorithm (MC-Watershed), and Split and Merge Watershed (SM-Watershed) [16].

The DSC and IoU metrics reported in Table 3 are computed against the gold standard automatically obtained by processing the DAPI channel. To obtain a fair comparison with the results presented in [16], we processed the 40 cell images from the homogeneous class used in that work. Even if the SNPHEp-2 dataset consists of cell images with different characteristics with respect to those contained in the VU and 2018 DSB datasets, ACDC achieved better results than CellProfiler, and slightly worse results than MC-Watershed and SM-Watershed approaches, which were specifically tailored to analyze this kind of images (see Table 3). These results were achieved by using the same settings in the pipeline, further showing that ACDC is a reliable method that can also be easily tuned to obtain satisfactory segmentation and cell counting outcomes on different datasets.

**Table 3.**
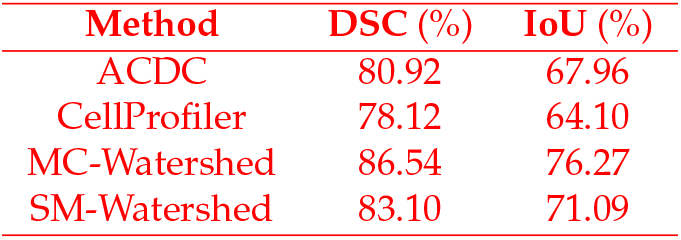
Comparison of ACDC, CellProfiler, MC-Watershed, and SM-Watershed in terms of cell nuclei segmentation on the SNPHEp-2 dataset.

## 4. Discussion and Conclusions

The fully automatic pipeline for cell detection and counting proposed in this paper, called ACDC, exploits watershed transform [39,40] and morphological filtering operations [41,42], and also benefits from the edge-preserving smoothing achieved by the bilateral filtering [38]. Notably, this pipeline does not depend on expert knowledge and can be deployed by setting only a few parameters. The capabilities of ACDC were tested on two different cell imaging datasets characterized by significantly different acquisition devices, specimens, image characteristics, and experimental conditions. ACDC was shown to be accurate and reliable in terms of cell counting and segmentation accuracy, thus representing a laboratory feasible solution also thanks to its computational efficiency. As a matter of fact, the Pearson coefficient achieved both in the case of the VU and DSB datasets was higher than 0.96, demonstrating an excellent agreement between automated cell count achieved with ACDC and the manual gold standard. The performance of ACDC were also compared with two pipelines defined with ImageJ; our results highlighted that while the cell counting was comparable, ACDC allowed us to achieved better segmentation outcomes. Moreover, we considered an additional cell images dataset and compared the outcome achieved with ACDC against CellProfiler, Marker-Controlled Watershed and Split and Merge Watershed, further showing the reliability of ACDC for what concerns segmentation and cell counting results. Finally, with the current ACDC implementation, it is also possible to distribute the computation on multi-core architectures and computer clusters, to further reduce the running time required to analyze large single image stacks. To this end, an asynchronous job queue, based on a distributed message passing paradigm was developed exploiting Celery and RabbitMQ.

The time-lapse microscopy image samples analyzed in this study are considerably different from to the publicly available microscopy images of cell nuclei, such as in [76,77] and [75]. Indeed, these datasets were captured under highly different microscope configurations compared to the VU experiments; with particular interest to magnification, a 40 × /0.9 numerical aperture and a 20 × /0.8 objective lens were used in [76,77] and [75], respectively. These acquisition characteristics, summarized in Table 4, produce substantially different images, compared to the datasets analyzed in this work, as in the case of the average coverage (i.e., the percentage of pixels covered by cell nuclei), which is remarkably different. As a future development, we plan to improve ACDC, to achieve accurate cell nuclei detection and analysis for different cell imaging scenarios, independently from the acquisition methods.

**Table 4.** Main characteristics of fluorescence microscopy datasets.

|  | U2OS [76] | NIH3T3 [77] | VU | DSB [37,46] |
| --- | --- | --- | --- | --- |
| # images | 48 | 49 | 46 | 301 |
| Image size | $1349 \times 1030$ | $1344 \times 1024$ | $2160 \times 2160$ | various |
| Magnification | $40\times$ | $40\times$ | $10\times$ | various |
| Total #cells | 1831 | 2178 | 14045 | 7931 |
| Min. #cells | 24 | 29 | 22 | 1 |
| Max. #cells | 63 | 70 | 1763 | 124 |
| Avg. %coverage | 23% | 18% | 2.07% | 5.41% |

We also aim at exploiting the most recent Machine Learning techniques [78] to accurately refine the segmentation results. The improvements may be achieved by classifying the geometrical and textural features extracted from the detected cells [79]. This advanced computational analysis can allow us to gain biological insights into complex cellular processes [17]. Finally, as a biological application, in the near future we plan to focus on the Fluorescent, Ubiquitination-based Cell-Cycle Indicator (FUCCI) reporter system for investigating cell-cycle states [45], by combining accurate nuclei segmentation results, on multiple fluorescent proteins for cell labeling [80], and live-cell imaging of cell cycle and division.

## Author Contributions

Conceptualization, L.R., A.T., D.R.T., C.F.L., and P.C.; Investigation, L.R., A.T., D.R.T., D.B., V.Q., C.F.L., and P.C.; Methodology, L.R., A.T., D.R.T., C.F.L., and P.C.; Software, L.R., A.T., D.R.T., and A.L.R.L.; Validation, L.R., A.T., D.R.T., R.B., C.M., C.F.L., and P.C.; Writing – original draft, L.R., A.T., D.R.T., A.L.R.L., C.F.L., and P.C.; Writing – review & editing, R.B., C.M., S.S., M.S.N., D.B., A.L.R.L., V.Q., and G.M.; Visualization, L.R., A.T., D.R.T., C.F.L., and P.C.; Supervision, D.R.T., G.M., C.F.L., and P.C.. All authors read and approved the final manuscript.

## Acknowledgments

The authors would like to thank Prof. Margarita Gamarra for her help with the analysis performed on the SNP HEp-2 Cell Dataset.

## References

1. Kanade, T.; Yin, Z.; Bise, R.; Huh, S.; Eom, S.; Sandbothe, M.F.; Chen, M. Cell image analysis: Algorithms, system and applications. Proceedings of the IEEE Workshop on Applications of Computer Vision (WACV). IEEE, 2011, pp. 374–381. doi:10.1109/WACV.2011.5711528.

2. Orth, J.D.; Kohler, R.H.; Foijer, F.; Sorger, P.K.; Weissleder, R.; Mitchison, T.J. Analysis of mitosis and antimitotic drug responses in tumors by in vivo microscopy and single-cell pharmacodynamics. Cancer Res. 2011, 71, 4608–4616. doi:10.1158/0008-5472.CAN-11-0412.

3. Manandhar, S.; Bouthemy, P.; Welf, E.; Danuser, G.; Roudot, P.; Kervrann, C. 3D flow field estimation and assessment for live cell fluorescence microscopy. Bioinformatics 2020, 36, 1317–1325. doi:10.1093/bioinformatics/btz780.

4. Peng, H. Bioimage informatics: a new area of engineering biology. Bioinformatics 2008, 24, 1827–1836. doi:10.1093/bioinformatics/btn346.

5. Meijering, E.; Carpenter, A.E.; Peng, H.; Hamprecht, F.A.; Olivo-Marin, J.C. Imagining the future of bioimage analysis. Nat. Biotechnol. 2016, 34, 1250. doi:10.1038/nbt.3722.

6. Peng, H.; Bateman, A.; Valencia, A.; Wren, J.D. Bioimage informatics: a new category in Bioinformatics. Bioinformatics 2012,28, 1057–1057. doi:10.1093/bioinformatics/bts111.

7. Meijering, E. Cell segmentation: 50 years down the road [life sciences]. IEEE Signal Process. Mag. 2012, 29,140–145. doi:10.1109/MSP.2012.2204190.

8. Schneider, C.A.; Rasband, W.S.; Eliceiri, K.W. NIH Image to ImageJ: 25 years of image analysis. Nat. Methods 2012, 9, 671. doi:10.1038/nmeth.2089.

9. Schindelin, J.; Arganda-Carreras, I.; Frise, E.; Kaynig, V.; Longair, M.; Pietzsch, T.; Preibisch, S.; Rueden, C.; Saalfeld, S.; Schmid, B.; others. Fiji: an open-source platform for biological-image analysis. Nat. Methods 2012, 9, 676. doi:10.1038/nmeth.2019.

10. Carpenter, A.E.; Jones, T.R.; Lamprecht, M.R.; Clarke, C.; Kang, I.H.; Friman, O.; others. CellProfiler: image analysis software for identifying and quantifying cell phenotypes. Genome Biol. 2006, 7, R100. doi:10.1186/gb-2006-7-10-r100.

11. Dao, D.; Fraser, A.N.; Hung, J.; Ljosa, V.; Singh, S.; Carpenter, A.E. CellProfiler Analyst: interactive data exploration, analysis and classification of large biological image sets. Bioinformatics 2016, 32, 3210–3212. doi:10.1093/bioinformatics/btw390.

12. Wählby, C.; Sintorn, I.M.; Erlandsson, F.; Borgefors, G.; Bengtsson, E. Combining intensity, edge and shape information for 2D and 3D segmentation of cell nuclei in tissue sections. J. Microsc. 2004, 215, 67–76. doi:10.1111/j.0022-2720.2004.01338.x.

13. Kaliman, S.; Jayachandran, C.; Rehfeldt, F.; Smith, A.S. Limits of Applicability of the Voronoi Tessellation Determined by Centers of Cell Nuclei to Epithelium Morphology. Front. Physiol. 2016, 7, 551. doi:10.3389/fphys.2016.00551.

14. Honda, H. Description of cellular patterns by Dirichlet domains: the two-dimensional case. J. Theor. Biol. 1978, 72, 523–543. doi:10.1016/0022-5193(78)90315-6.

15. Kostrykin, L.; Schnörr, C.; Rohr, K. Globally optimal segmentation of cell nuclei in fluorescence microscopy images using shape and intensity information. Med. Image Anal. 2019, 58, 101536. doi:10.1016/j.media.2019.101536.

16. Gamarra, M.; Zurek, E.; Escalante, H.J.; Hurtado, L.; San-Juan-Vergara, H. Split and merge watershed: a two-step method for cell segmentation in fluorescence microscopy images. Biomed. Signal Process. Control 2019, 53, 101575. doi:10.1016/j.bspc.2019.101575.

17. Angermueller, C.; Pärnamaa, T.; Parts, L.; Stegle, O. Deep learning for computational biology. Mol. Syst. Biol. 2016, 12, 878. doi:10.15252/msb.20156651.

18. Berg, S.; Kutra, D.; Kroeger, T.; Straehle, C.N.; Kausler, B.X.; Haubold, C.; others. ilastik: interactive machine learning for (bio) image analysis. Nat. Methods 2019,16, 1226–1232. doi:10.1038/s41592-019-0582-9.

19. Held, M.; Schmitz, M.H.; Fischer, B.; Walter, T.; Neumann, B.; Olma, M.H.; Peter, M.; Ellenberg, J.; Gerlich, D.W. CellCognition: time-resolved phenotype annotation in high-throughput live cell imaging. Nat. Methods 2010, 7, 747. doi:10.1038/nmeth.1486.

20. Ciresan, D.; Giusti, A.; Gambardella, L.M.; Schmidhuber, J. Deep neural networks segment neuronal membranes in electron microscopy images. Advances in Neural Information Processing Systems (NIPS), 2012, pp. 2843–2851.

21. Rosati, R.; Romeo, L.; Silvestri, S.; Marcheggiani, F.; Tiano, L.; Frontoni, E. Faster R-CNN approach for detection and quantification of DNA damage in comet assay images. Comput. Biol. Med. 2020, p. 103912. doi:10.1016/j.compbiomed.2020.103912.

22. Sadanandan, S.K.; Ranefall, P.; Le Guyader, S.; Wählby, C. Automated Training of Deep Convolutional Neural Networks for Cell Segmentation. Sci. Rep. 2017, 7, 7860. doi:10.1038/s41598-017-07599-6.

23. Hiramatsu, Y.; Hotta, K.; Imanishi, A.; Matsuda, M.; Terai, K.; Liu, D.; Zhang, D.; Song, Y.; Zhang, C.; Huang, H.; others. Cell Image Segmentation by Integrating Multiple CNNs. Proceedings of the IEEE Conference on Computer Vision and Pattern Recognition (CVPR) Workshops, 2018, pp. 2205–2211.

24. Ren, S.; He, K.; Girshick, R.; Sun, J. Faster R-CNN: Towards real-time object detection with region proposal networks. Proceedings of the Advances in Neural Information Processing Systems (NIPS), 2015, pp. 91–99.

25. Redmon, J.; Divvala, S.; Girshick, R.; Farhadi, A. You only look once: Unified, real-time object detection. Proceedings of the Conference on Computer Vision and Pattern Recognition (CVPR). IEEE, 2016, pp. 779–788. doi:10.1109/CVPR.2016.91.

26. Alam, M.M.; Islam, M.T. Machine learning approach of automatic identification and counting of blood cells. Healthc. Technol. Lett. 2019, 6, 103–108. doi:10.1049/htl.2018.5098.

27. Han, C.; Kitamura, Y.; Kudo, A.; others. Synthesizing diverse lung nodules wherever massively: 3D multi-conditional GAN-based CT image augmentation for object detection. Proceedings of the International Conference on 3D Vision (3DV). IEEE, 2019, pp. 729–737. doi:10.1109/3DV.2019.00085.

28. Bayramoglu, N.; Heikkilä, J. Transfer learning for cell nuclei classification in histopathology images. Proceedings of the European Conference on Computer Vision (ECCV) Workshops. Springer, 2016, Vol. 9915, *LNCS*, pp. 532–539. doi:10.1007/978-3-319-49409-8\_46.

29. Apicella, A.; Isgrò, F.; Prevete, R. A simple and efficient architecture for trainable activation functions. Neurocomputing 2019, 370, 1–15. doi:10.1016/j.neucom.2019.08.065.

30. Pelt, D.M.; Sethian, J.A. A mixed-scale dense convolutional neural network for image analysis. Proc. Natl. Acad. Sci. 2018,115, 254–259. doi:10.1073/pnas.1715832114.

31. Osokin, A.; Chessel, A.; Carazo Salas, R.E.; Vaggi, F. GANs for biological image synthesis. Proceedings of the IEEE International Conference on Computer Vision (ICCV). IEEE, 2017, pp. 2233–2242. doi:10.1109/ICCV.2017.245.

32. Han, C.; Rundo, L.; Araki, R.; Furukawa, Y.; Mauri, G.; Nakayama, H.; Hayashi, H. Infinite brain MR images: PGGAN-based data augmentation for tumor detection. In Neural Approaches to Dynamics of Signal Exchanges; Springer, 2019; Vol. 151, *Smart Innovation, Systems and Technologies*, pp. 291–303. doi:10.1007/978-981-13-8950-4_27.

33. Lo Castro, D.; Tegolo, D.; Valenti, C. A visual framework to create photorealistic retinal vessels for diagnosis purposes. J. Biomed. Inform. 2020, p. 103490. doi:10.1016/j.jbi.2020.103490.

34. Kraus, O.Z.; Ba, J.L.; Frey, B.J. Classifying and segmenting microscopy images with deep multiple instance learning. Bioinformatics 2016, 32, i52–i59. doi:10.1093/bioinformatics/btw252.

35. Militello, C.; Rundo, L.; Minafra, L.; Cammarata, F.P.; Calvaruso, M.; Conti, V.; Russo, G. MF2C3: Multi-Feature Fuzzy Clustering to Enhance Cell Colony Detection in Automated Clonogenic Assay Evaluation. Symmetry 2020, 12, 773. doi:10.3390/sym12050773.

36. Meyer, C.T.; Wooten, D.J.; Paudel, B.B.; Bauer, J.; Hardeman, K.N.; Westover, D.; Lovly, C.M.; Harris, L.A.; Tyson, D.R.; Quaranta, V. Quantifying drug combination synergy along potency and efficacy axes. Cell Syst. 2019, 8, 97–108. doi:10.1016/j.cels.2019.01.003.

37. Caicedo, J.C.; Goodman, A.; Karhohs, K.W.; Cimini, B.A.; Ackerman, J.; Haghighi, M.; others. Nucleus segmentation across imaging experiments: the 2018 Data Science Bowl. Nat. Methods 2019, 16, 1247–1253. doi:10.1038/s41592-019-0612-7.

38. Tomasi, C.; Manduchi, R. Bilateral filtering for gray and color images. Proceedings of the Sixth International Conference on Computer Vision (ICCV). IEEE, 1998, pp. 839–846. doi:10.1109/ICCV.1998.710815.

39. Soille, P.J.; Ansoult, M.M. Automated basin delineation from digital elevation models using mathematical morphology. Signal Process. 1990, 20, 171–182. doi:10.1016/0165-1684(90)90127-K.

40. Vincent, L.; Soille, P. Watersheds in digital spaces: an efficient algorithm based on immersion simulations. IEEE Trans. Pattern Anal. Mach. Intell. 1991, 13, 583–598. doi:10.1109/34.87344.

41. Beucher, S.; Meyer, F. The morphological approach to segmentation: the watershed transformation. In Mathematical Morphology in Image Processing; Marcel Dekker Inc.: New York. NY, USA, 1993; Vol. 34, pp. 433–481.

42. Soille, P. Morphological Image Analysis: Principles and Applications, 2 ed.; Springer Science & Business Media: Secaucus, NJ, USA, 2004. doi:10.1007/978-3-662-05088-0.

43. Tyson, D.R.; Garbett, S.P.; Frick, P.L.; Quaranta, V. Fractional proliferation: a method to deconvolve cell population dynamics from single-cell data. Nat. Methods 2012, 9, 923. doi:10.1038/nmeth.2138.

44. Harris, Leonard A.; Frick, Peter L.; Garbett, Shawn P.; Hardeman, Keisha N.; Paudel, B Bishal.; Lopez, Carlos F.; Quaranta, Vito.; Tyson, Darren R. An unbiased metric of antiproliferative drug effect in vitro. Nat. Methods 2016, 13, 497–500. doi:10.1038/nmeth.3852.

45. Sakaue-Sawano, A.; Kurokawa, H.; Morimura, T.; Hanyu, A.; Hama, H.; Osawa, H.; Kashiwagi, S.; Fukami, K.; Miyata, T.; Miyoshi, H.; others. Visualizing spatiotemporal dynamics of multicellular cell-cycle progression. Cell 2008, 132, 487–498. doi:10.1016/j.cell.2007.12.033.

46. Kaggle. 2018 Data Science Bowl. https://www.kaggle.com/c/data-science-bowl-2018, 2018. Online; accessed 14 December 2019.

47. Georgescu, W.; Wikswo, J.P.; Quaranta, V. CellAnimation: an open source MATLAB framework for microscopy assays. Bioinformatics 2011, 28, 138–139. doi:10.1093/bioinformatics/btr633.

48. Sansone, M.; Zeni, O.; Esposito, G. Automated segmentation of comet assay images using Gaussian filtering and fuzzy clustering. Med. Biol. Eng. Comput. 2012, 50, 523–532. doi:10.1007/s11517-012-0882-z.

49. Schettini, R.; Gasparini, F.; Corchs, S.; Marini, F.; Capra, A.; Castorina, A. Contrast image correction method. J. Electron. Imaging 2010, 19, 023005. doi:10.1117/1.3386681.

50. Venkatesh, M.; Mohan, K.; Seelamantula, C.S. Directional bilateral filters for smoothing fluorescence microscopy images. AIP Advances 2015, 5, 084805. doi:10.1063/1.4930029.

51. Jiang, W.; Baker, M.L.; Wu, Q.; Bajaj, C.; Chiu, W. Applications of a bilateral denoising filter in biological electron microscopy. J. Struct. Biol. 2003, 144, 114–122. doi:10.1016/j.jsb.2003.09.028.

52. Li, K.; Miller, E.D.; Chen, M.; Kanade, T.; Weiss, L.E.; Campbell, P.G. Computer vision tracking of stemness. Proceedings of the 5th IEEE International Symposium on Biomedical Imaging: From Nano to Macro (ISBI). IEEE, 2008, pp. 847–850. doi:10.1109/ISBI.2008.4541129.

53. Gonzalez, R.; Woods, R. Digital Image Processing, 3 ed.; Prentice Hall Press: Upper Saddle River, NJ, USA, 2002.

54. Jain, A.K. Fundamentals of Digital Image Processing, 1 ed.; Prentice Hall Press: Upper Saddle River, NJ, USA, 2002.

55. Otsu, N. A threshold selection method from gray-level histograms. IEEE Trans. Syst. Man Cybern. 1975, 11, 23–27. doi:10.1109/TSMC.1979.4310076.

56. Militello, C.; Rundo, L.; Conti, V.; Minafra, L.; Cammarata, F.P.; Mauri, G.; Gilardi, M.C.; Porcino, N. Area-based cell colony surviving fraction evaluation: A novel fully automatic approach using general-purpose acquisition hardware. Comput. Biol. Med. 2017, 89, 454–465. doi:10.1016/j.compbiomed.2017.08.005.

57. Borgefors, G. Distance transformations in digital images. Comput. Vis. Graph. Image Process. 1986, 34, 344–371. doi:10.1016/S0734-189X(86)80047-0.

58. Felzenszwalb, P.F.; Huttenlocher, D.P. Distance transforms of sampled functions. Theory Comput. 2012, 8, 415–428. doi:10.4086/toc.2012.v008a019.

59. Salvi, M.; Morbiducci, U.; Amadeo, F.; Santoro, R.; Angelini, F.; Chimenti, I.; Massai, D.; Messina, E.; Giacomello, A.; Pesce, M.; others. Automated segmentation of fluorescence microscopy images for 3D cell detection in human-derived cardiospheres. Sci. Rep. 2019, 9, 6644. doi:10.1038/s41598-019-43137-2.

60. Grau, V.; Mewes, A.; Alcaniz, M.; Kikinis, R.; Warfield, S.K. Improved watershed transform for medical image segmentation using prior information. IEEE Trans. Med. Imaging 2004, 23, 447–458. doi:10.1109/TMI.2004.824224.

61. Suzuki, K.; Horiba, I.; Sugie, N. Linear-time connected-component labeling based on sequential local operations. Comput. Vis. Image Underst. 2003, 89, 1–23. doi:10.1016/S1077-3142(02)00030-9.

62. Meyer, F. Topographic distance and watershed lines. Signal Process. 1994, 38, 113–125. doi:10.1016/0165-1684(94)90060-4.

63. Najman, L.; Couprie, M.; Bertrand, G. Watersheds, mosaics, and the emergence paradigm. Discrete Appl. Math. 2005, 147, 301–324. doi:10.1016/j.dam.2004.09.017.

64. van der Walt, S.; Schönberger, J.L.; Nunez-Iglesias, J.; Boulogne, F.; Warner, J.D.; Yager, N.; Gouillart, E.; Yu, T.; the scikit-image contributors. scikit-image: image processing in Python. PeerJ 2014, 2, e453. doi:10.7717/peerj.453.

65. Coelho, L.P. Mahotas: Open source software for scriptable computer vision. J. Open Res. Softw. 2013, 1, e3. doi:10.5334/jors.ac.

66. Rundo, L.; Militello, C.; Vitabile, S.; Casarino, C.; Russo, G.; Midiri, M.; Gilardi, M.C. Combining split-and-merge and multi-seed region growing algorithms for uterine fibroid segmentation in MRgFUS treatments. Med. Biol. Eng. Comput. 2016, 54, 1071–1084. doi:10.1007/s11517-015-1404-6.

67. Celery Project. Celery Distributed Task Queue. http://www.celeryproject.org/, 2018. Online; accessed 14 December 2019.

68. Pivotal Software, Inc. RabbitMQ. http://www.rabbitmq.com/, 2018. Online; accessed 14 December 2019.

69. Dalcín, L.; Paz, R.; Storti, M. MPI for Python. J. Parallel Distrib. Comput. 2005, 65, 1108–1115. doi:10.1016/j.jpdc.2005.03.010.

70. Rundo, L.; Tangherloni, A.; Cazzaniga, P.; Nobile, M.S.; Russo, G.; Gilardi, M.C.; others. A novel framework for MR image segmentation and quantification by using MedGA. Comput. Methods Programs Biomed. 2019, 176, 159–172. doi:10.1016/j.cmpb.2019.04.016.

71. Tangherloni, A.; Spolaor, S.; Rundo, L.; Nobile, M.S.; Cazzaniga, P.; Mauri, G.; Liò, P.; Merelli, I.; Besozzi, D. GenHap: a novel computational method based on genetic algorithms for haplotype assembly. BMC Bioinformatics 2019, 20, 172. doi:10.1186/s12859-019-2691-y.

72. Tangherloni, A.; Rundo, L.; Spolaor, S.; Cazzaniga, P.; Nobile, M.S. GPU-powered multi-swarm parameter estimation of biological systems: A master-slave approach. Proceedings of the 26th Euromicro International Conference on Parallel, Distributed and Network-based Processing (PDP). IEEE, 2018, pp. 698–705.

73. Bland, J.M.; Altman, D.G. Measuring agreement in method comparison studies. Stat. Methods Med. Res. 1999, 8, 135–160. doi:10.1177/096228029900800204.

74. Wilcoxon, F. Individual comparisons by ranking methods. Biometrics Bull. 80–83, 1, 196–202. doi:10.2307/3001968.

75. Wiliem, A.; Wong, Y.; Sanderson, C.; Hobson, P.; Chen, S.; Lovell, B.C. Classification of human epithelial type 2 cell indirect immunofluoresence images via codebook based descriptors. Proceedings of the IEEE Workshop on Applications of Computer Vision (WACV). IEEE, 2013, pp. 95–102. doi:10.1109/WACV.2013.6475005.

76. Coelho, L.P.; Shariff, A.; Murphy, R.F. Nuclear segmentation in microscope cell images: a hand-segmented dataset and comparison of algorithms. Proceedings of the IEEE International Symposium on Biomedical Imaging (ISBI): From Nano to Macro. IEEE, 2009, pp. 518–521. doi:10.1109/ISBI.2009.5193098.

77. Osuna, E.G.; Hua, J.; Bateman, N.W.; Zhao, T.; Berget, P.B.; Murphy, R.F. Large-scale automated analysis of location patterns in randomly tagged 3T3 cells. Ann. Biomed. Eng. 2007, 35, 1081–1087. doi:10.1007/s10439-007-9254-5.

78. Kraus, O.Z.; Grys, B.T.; Ba, J.; Chong, Y.; Frey, B.J.; Boone, C.; Andrews, B.J. Automated analysis of high-content microscopy data with deep learning. Mol. Syst. Biol. 2017, 13, 924. doi:10.15252/msb.20177551.

79. Win, K.; Choomchuay, S.; Hamamoto, K.; Raveesunthornkiat, M. Detection and Classification of Overlapping Cell Nuclei in Cytology Effusion Images Using a Double-Strategy Random Forest. Appl. Sci. 2018, 8, 1608. doi:10.3390/app8091608.

80. Salvi, M.; Cerrato, V.; Buffo, A.; Molinari, F. Automated segmentation of brain cells for clonal analyses in fluorescence microscopy images. J. Neurosci. Methods 2019, 325, 108348. doi:10.1016/j.jneumeth.2019.108348.

